# Reengineering mRNA lipid nanoparticles for systemic delivery to pancreas

**DOI:** 10.1101/2024.10.30.621163

**Authors:** Ivan Isaac, Luv Patel, Nguyen Tran, Amarnath Singam, Prasun Guha, Seungman Park, Chandrabali Bhattacharya

## Abstract

Lipid nanoparticles (LNPs) hold transformative potential for nucleic acid delivery, with applications ranging from clinical use, particularly in COVID-19 vaccines, to gene therapy and cancer immunotherapy. Traditional LNPs are composed of four components: ionizable lipids, cholesterol, helper lipids, and PEGylated lipids. However, a significant hurdle remains the need for more efficient and selective systemic delivery vehicles, as most targets are challenging to reach and primarily accumulate in the liver following intravenous administration, largely due to Apolipoprotein E (ApoE) mediated uptake in the blood. Recent studies have shown that introducing cationic or anionic lipids as a fifth component in the LNPs has resulted in lung- and spleen-specific mRNA expression. In this study, we report that incorporating endogenous ligands, such as vitamins, as a fifth component can enhance the extrahepatic localization of LNPs. Vitamins are highly biocompatible, possess excellent targeting potential, influence immune response, improve cellular uptake, and increase the stability of mRNA LNPs. We developed a library of 100 LNPs containing luciferase mRNA by partially replacing ionizable lipids with this fifth component and evaluated their efficacy both in vitro and in vivo. From comprehensive batch analysis screening, we identified two formulations with cholecalciferol (vitamin D3) as a fifth component that demonstrated selective systemic delivery to the pancreas (> 99%) with high efficacy. Among these, C-CholF3 emerged as the top formulation, exhibiting robust and sustained protein expression in the pancreas for up to 3 days in a dose-dependent manner with minimal toxicity that makes it suitable for repeated administration. Furthermore, C-CholF3 also demonstrated pancreas-specific gene editing in the Ai14 transgenic mouse model, showing high expression of TdTomato. This underscores its translational potential for protein replacement and CRISPR/Cas9-mediated gene editing in currently incurable pancreatic diseases.

## Introduction

Lipid nanoparticles (LNPs) have revolutionized the field of nucleic acid delivery and have been extensively explored in gene therapy, protein replacement therapies, and immunotherapies for various diseases.^1–3^ Their clinical applications were notably demonstrated by their widespread use in delivering the SARS-CoV-2 vaccine during the COVID-19 pandemic.^4–7^ Despite their promise, targeting beyond the liver remains a significant challenge with the traditional LNPs, which are made up of four key components —ionizable lipid, phospholipid, cholesterol, and polyethylene glycol (PEG) lipid.^[5,8,9]^ The accumulation of LNPs in the liver primarily occurs because they associate with Apolipoprotein E (ApoE) in the blood, which is crucial for cholesterol transport and low-density lipoprotein receptor (LDLR)-mediated endocytosis.^10–13^ One promising approach for redirecting LNPs to extrahepatic sites is incorporating a fifth component in the traditional LNP formulation. This strategy has been effective in achieving lung- and spleen-specific mRNA expression through interactions with plasma proteins, such as vitronectin (Vtn) and β2 glycoprotein 1 (β2-GPI), using SORT LNPs containing a fifth component cationic or anionic lipid.^14–18^ Furthermore, the addition of an adjuvant lipidoid containing a Toll-like receptor (TLR) 7/8 agonist as a fifth component also successfully demonstrated transfection in the lymph nodes.^[19]^ Peptides such as MH42 and RGD were utilized to design targeted LNPs for the retina and bone, employing a fifth-component system as well.^[20,21]^ This illustrates the potential of LNPs to target other hard-to-reach tissues with formulation design, overcoming the inherent limitations of classical four-component LNPs.

Here, we report that adding endogenous ligands such as vitamins as the fifth component in the LNP formulation could fine-tune their organ tropism and enhance targeted delivery (**Figure 1A**). Their distinct chemical structures and unique biochemical functions make them ideal candidates for the fifth component platform.^22–24^ Structurally, vitamins can be broadly divided into water-soluble groups, such as the vitamin B complex, and fat-soluble groups, including vitamins A, D, E, and K.^[25,26]^ The fat-soluble vitamins are particularly relevant to the LNP formulations due to their hydrophobic tails, which can integrate seamlessly into the lipid bilayer.^[27]^ The long hydrophobic isoprenoid chain in vitamin A may facilitate membrane fusion and cellular uptake. ^[33]^ Vitamin D, being a secosteroid, binds to vitamin D receptors (VDRs) expressed in specific tissues, including the pancreas. Similarly, vitamin E is also a very potent antioxidant with a hydrophobic chromanol ring and phytyl tail, which may stabilize the lipid nanoparticles, reduce oxidative damage, and redirect biodistribution to extrahepatic tissues. In this study, we screened a library of 100 LNPs formulated with different vitamins as a fifth component, combined with five different ionizable lipids and four different formulation ratios, to evaluate their potential for tissue-specific delivery (**Figure 1B**). The LNPs were tested in five different cell lines representing major cell types in the body, and we observed varying levels of transfection across these cell types. We further investigated the ability of these LNPs to deliver mRNA in vivo using batch analysis.

**Figure 1.**
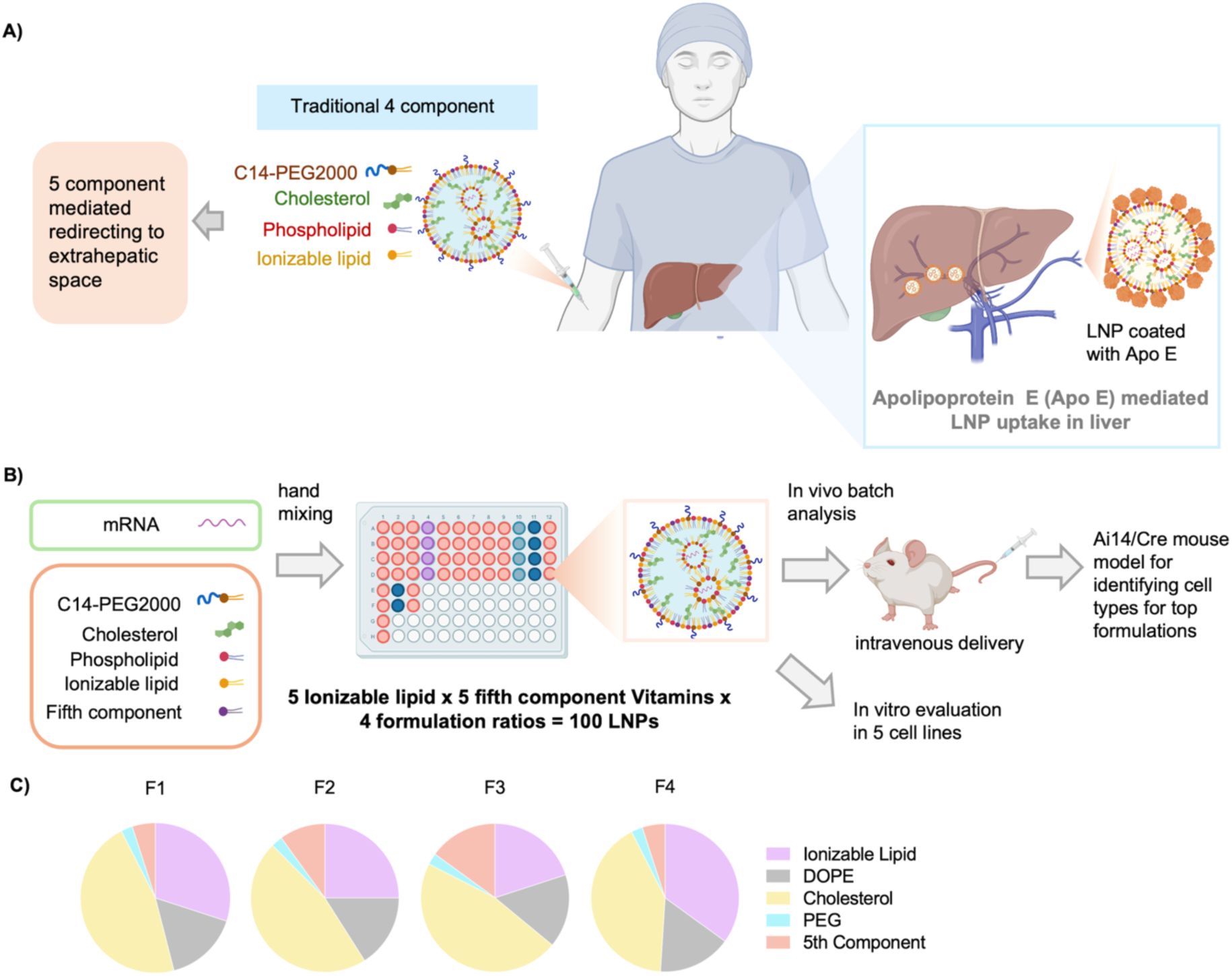
Overview of the fifth component-based mRNA LNP formulation and screening platform. **A)** Schematic representation illustrating the current challenges in developing a robust extrahepatic delivery system. **B)** Schematic showing our fifth component-based mRNA LNP formulation and screening platform. **C)** Pie charts depicting the different formulation ratios for ionizable lipid, phospholipid, cholesterol, PEG lipid, and endogenous vitamin fifth component.

This screening led to the identification of two formulations containing cholecalciferol as a fifth component, which demonstrated selective delivery of mRNA to the pancreas with high efficacy through intravenous administration - C-CholF2 and C-CholF3. Among these formulations, C-CholF3 exhibited higher protein expression than C-CholF2, achieving over 99% selectivity and sustained expression in the pancreas for up to 3 days in a dose-dependent manner. The addition of biocompatible vitamins also resulted in decreased toxicity, resulting in a better safety profile and improved tolerance. Furthermore, we assessed the gene editing ability of C-CholF3 LNP in the Ai14 mouse model and found tissue-specific gene editing in the pancreas, with efficient expression of TdTomato. In summary, this platform highlights the importance of incorporating vitamins as the fifth component of LNPs for targeted delivery to specific tissues. Furthermore, it broadens the potential for targeting hard-to-reach extrahepatic organs, such as the pancreas, and provides solutions for protein replacement and CRISPR/Cas9-mediated gene editing to treat currently incurable pancreatic diseases.

## Results and Discussion

Traditional LNPs for mRNA delivery consist of four key components, each tailored to establish a stable lipid bilayer structure. These lipids include: 1) ionizable lipids for complexing with mRNA with charge-altering properties, facilitating the release of mRNA cargo in the acidic endosomal environment; 2) helper lipids for bilayer structure reinforcement; 3) cholesterol for increasing structural integrity; and 4) PEG-lipids for extending circulation in the bloodstream.^28–31^ To enhance organ-specific delivery of LNPs, we added vitamins as a fifth component in the formulation. The vitamins A, B2, D3, E, or K1 were chosen for their unique biological functions, which can complement the LNP core in different ways. Vitamin A (retinol) is well-known for its role in cell differentiation and immune modulation and, could enhance the targeting to immune cells and tissues that display high retinoid activity.^[32,33]^ The lipophilic nature of this molecule allows for easy insertion into the lipid bilayer and may facilitate membrane fusion during its cellular uptake.^[34]^ Vitamin B2 (riboflavin), although water soluble, was included because it plays crucial role in cellular metabolism and energy production.^[35,36]^ Vitamin D3 (cholecalciferol) plays a key role in calcium homeostasis and immune function, with the potential to improve biodistribution in tissues expressing high levels of the Vitamin D receptor (VDR), notably the pancreas.^[37]^ This vitamin is essential for maintaining β-cell function, supporting insulin release, reducing β-cell apoptosis, and promoting overall cell health.^[38]^ Vitamin E (tocopherol) possesses significant antioxidant properties and could be incorporated to safeguard the lipid nanoparticles (LNPs) from oxidative damage during circulation, thereby prolonging their half-life and enhancing delivery to specific target cells.^[39]^ Furthermore, Vitamin K1 (phylloquinone) is crucial for blood coagulation and bone health and could target tissues associated with coagulation pathways.^[40,41]^

To further diversify our vitamin-based LNP design, we developed a comprehensive formulation library using previously studied ionizable lipids (ILs), including benchmark SM-102, MC3, and C12-200, which are known to traffic to the liver.^42–44^ Additionally, we used THP1, which our research group previously established as an effective mRNA delivery system.^[45]^ 306Oi10 was also included, as it showed promise in preclinical studies by exhibiting high transfection efficiency.^[46]^ For an efficient fifth-component formulation, it is crucial to partially replace the ionizable lipid without affecting encapsulation or endosomal escape. We started our study by using a benchmark formulation containing ionizable lipid, helper lipid, cholesterol, and DMG-PEG 2000 at a 35:16:46.5:2.5 molar ratio, which was then modified by partially replacing the ionizable lipid. The four different formulation ratios were chosen with an increasing vitamin substitution (5–15 molar ratio) to systematically investigate the effect of vitamin molar ratios on the performances of these LNPs. The detailed formulation ratios are shown in Figure 1C and **Table S1**. Designing the formulation ratio is as significant as selecting the suitable fifth component in this study as it impacts the physicochemical parameters of LNP, including particle size, encapsulation efficiency (EE%), and surface charge, which are important for efficient mRNA delivery. These interactions can often significantly change with minor changes in the lipid-to-vitamin ratio, altering the stability of particles in circulation and endosomal escape capability. We formulated a library of 100 different LNP formulations by combining five different ILs, five vitamins, and four different formulation ratios (5 ILs × 5 Vitamins × 4 Formulation = 100 LNPs) in a 96-well plate format using hand mixing. We used firefly luciferase (FLuc) mRNA as our reporter because its non-secretory nature allows direct imaging and quantification of transfection efficiency. The particle size and PDI of each LNP formulation were measured using dynamic light scattering (DLS), as shown in **Figures S1** and **S2**. The results indicate that the average diameter of most LNPs was within a window of 60 to 140 nm and PDIs smaller than 0.2, indicating a relatively narrow size distribution. The uniformity ensures efficient cellular uptake and successful endosomal escape, demonstrating that our fifth-component LNPs were structurally similar to conventional LNPs.

We assessed the in vitro transfection efficiencies of all the LNPs in various cell lines, including human embryonic kidney cells (HEK293), human foreskin fibroblasts (HFF), human umbilical vein endothelial cells (HUVEC), mouse macrophage cells (RAW264.7), and human microglia cells (HMC3). These cells represent a wide range of tissues such as the kidney, skin, vascular endothelium, immune system, and brain. A comprehensive analysis of the transfection data revealed several key trends, indicating a significant improvement in transfection by vitamin-based LNPs compared with control DLin-MC3-DMA (MC3) and SM-102. Notably, LNPs with riboflavin displayed significantly higher transfection in HEK293 cells compared with other vitamin-based formulations and the control LNPs (**Figure 2A**). This may be attributed to the crucial role played by riboflavin in cellular metabolism, energy generation, and ATP production.^[35,36,47]^ Incorporation of Riboflavin into LNPs may enhance metabolic pathways, promoting cellular uptake and LNP processing. LNPs formulated with retinol, riboflavin, and tocopherol, containing SM-102 or 306Oi10 ionizable lipids, showed significantly better transfection in HFF cells compared to controls and other formulations (**Figure 2B**). Fibroblast regeneration is crucial for skin regeneration and is greatly influenced by retinol due to its involvement in cellular differentiation.^[48]^ The improved transfection may be due to enhanced membrane fluidity and stabilizing the LNPs, thus allowing more efficient mRNA delivery in these cells. In HUVEC cells, which form the inner lining of the blood vessels, the LNPs incorporating riboflavin and tocopherol with C12-200 are among the highest transfecting formulations, outperforming controls (**Figure 2C**). Tocopherol helps protect endothelial cells from oxidative stress and promotes vascular health, which may improve mRNA delivery by increasing vascular permeability. ^[49]^ Additionally, the high transfection efficiency of riboflavin LNPs in HUVEC cells may also be connected to enhanced cellular metabolism.^[47]^ On the other hand, RAW264.7 cells, which resemble macrophages, exhibit significantly higher transfection with C12-200-based LNPs containing retinol (**Figure 2D**). This might be again attributed to the immunomodulatory effects of retinol that could improve cellular uptake and intracellular processing of LNPs.^50–52^ This suggests that retinol incorporation could improve immune cell targeting. In HMC3 microglial cells, we observed that LNPs containing retinol, riboflavin, and phylloquinone exhibited higher transfection (**Figure 2E**). Microglia are the resident immune cells of the brain. The receptors involved in the antioxidant and anti-inflammatory properties of these vitamins may promote receptor-mediated endocytosis and transfection.^53–56^ Hence, our findings demonstrate that incorporating the vitamins as a fifth component into the LNP formulations can greatly enhance transfection efficacy compared with traditional four-component LNPs in certain cell lines, providing a rationale for tissue-specific delivery.

**Figure 2.**
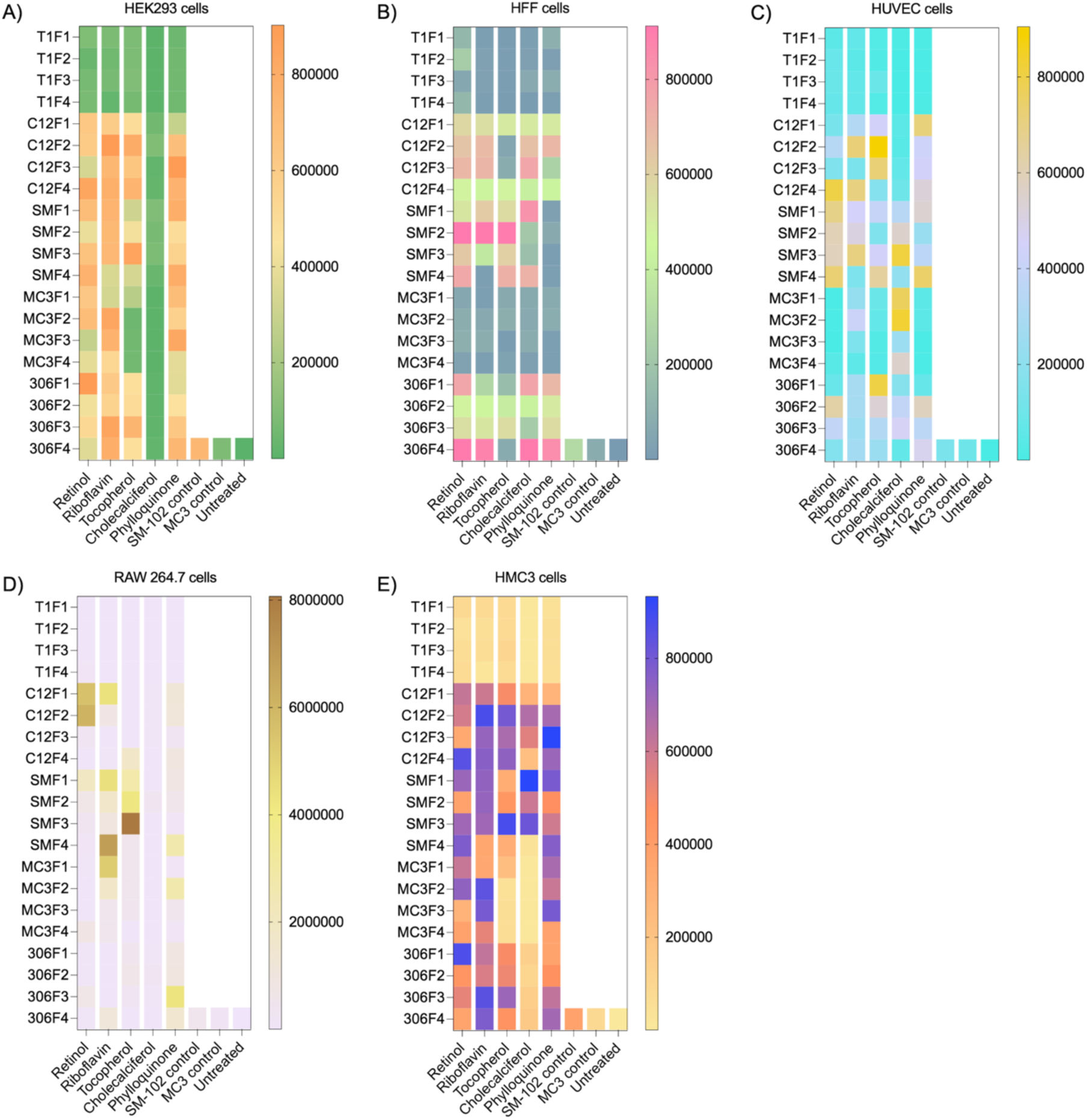
*In Vitro* screening of fifth component based LNPs. **A)** Heat map of in vitro delivery of luciferase mRNA with vitamins-based fifth component LNPs in HEK293 cells (125 ng mRNA per well, 96 well plate, n=3). Luminescence intensity was quantified 24 h after adding LNPs. **B)** Heat map of in vitro delivery of luciferase mRNA with vitamins-based fifth component LNPs in HFF cells (125 ng mRNA per well, 96 well plate, n=3). Luminescence intensity was quantified 24 h after adding LNPs. **C)** Heat map of in vitro delivery of luciferase mRNA with vitamins-based fifth component LNPs in HUVEC cells (125 ng mRNA per well, 96 well plate, n=3). Luminescence intensity was quantified 24 h after adding LNPs. **D)** Heat map of in vitro delivery of luciferase mRNA with vitamins-based fifth component LNPs in RAW 264.7 cells (125 ng mRNA per well, 96 well plate, n=3). Luminescence intensity was quantified 24 h after adding LNPs. **E)** Heat map of in vitro delivery of luciferase mRNA with vitamins-based fifth component LNPs in HMC3 cells (125 ng mRNA per well, 96 well plate, n=3). Luminescence intensity was quantified 24 h after adding LNPs.

Next, we evaluated the delivery efficacy of our fifth component, vitamin LNPs, in vivo using comprehensive batch analysis. This approach accelerated our screening process significantly and resulted in a remarkable reduction in the number of animals, as well as time and cost. In this study, we categorized 100 LNPs into four groups based on their formulation ratios (F1, F2, F3, and F4). Each group, containing 25 formulations corresponding to its respective formulation ratio, was administered to C57BL/6 mice via intravenous injection at a dosage of 0.5 mg/kg, and protein expression was assessed in major organs using an IVIS imaging system. Formulations F1 and F4 primarily showed transfection in the liver and, with additional expression in the spleen, intestines, and pancreas. Notably, formulations F2 and F3 demonstrated significant redirection to extrahepatic space, with F2 showing selective protein expression in the pancreas and minimal expression in other organs, including the spleen and liver (**Figure 3A**). More selective formulations, F2 and F3, were then batched based on ionizable lipids (THP1, C12-200, SM-102, MC3, and 306Oi10) in five groups containing both F2 and F3 LNPs, allowing for a total administration of 10 LNPs per injection at the same dose. Remarkably, C12-200 LNPs batch showed selective transfection in the pancreas, whereas the other ILs resulted in broader expression across various organs (**Figure 3B**). This highlights the significance of selecting the appropriate IL to achieve targeted mRNA delivery. Further, C12-200 was batched based on different vitamins (Retinol, Riboflavin, Cholecalciferol, Tocopherol, and Phylloquinone). Among these, cholecalciferol exhibited selective mRNA transfection and protein expression in the pancreas, reinforcing that the formulation ratios are critical in directing LNPs to the desired organ, highlighting the potential of vitamin-based LNPs for more precise mRNA delivery (**Figure 3C**). Throughout each phase of the screening process, SM-102 and MC3 were used as control LNPs with major accumulation in the liver.

**Figure 3.**
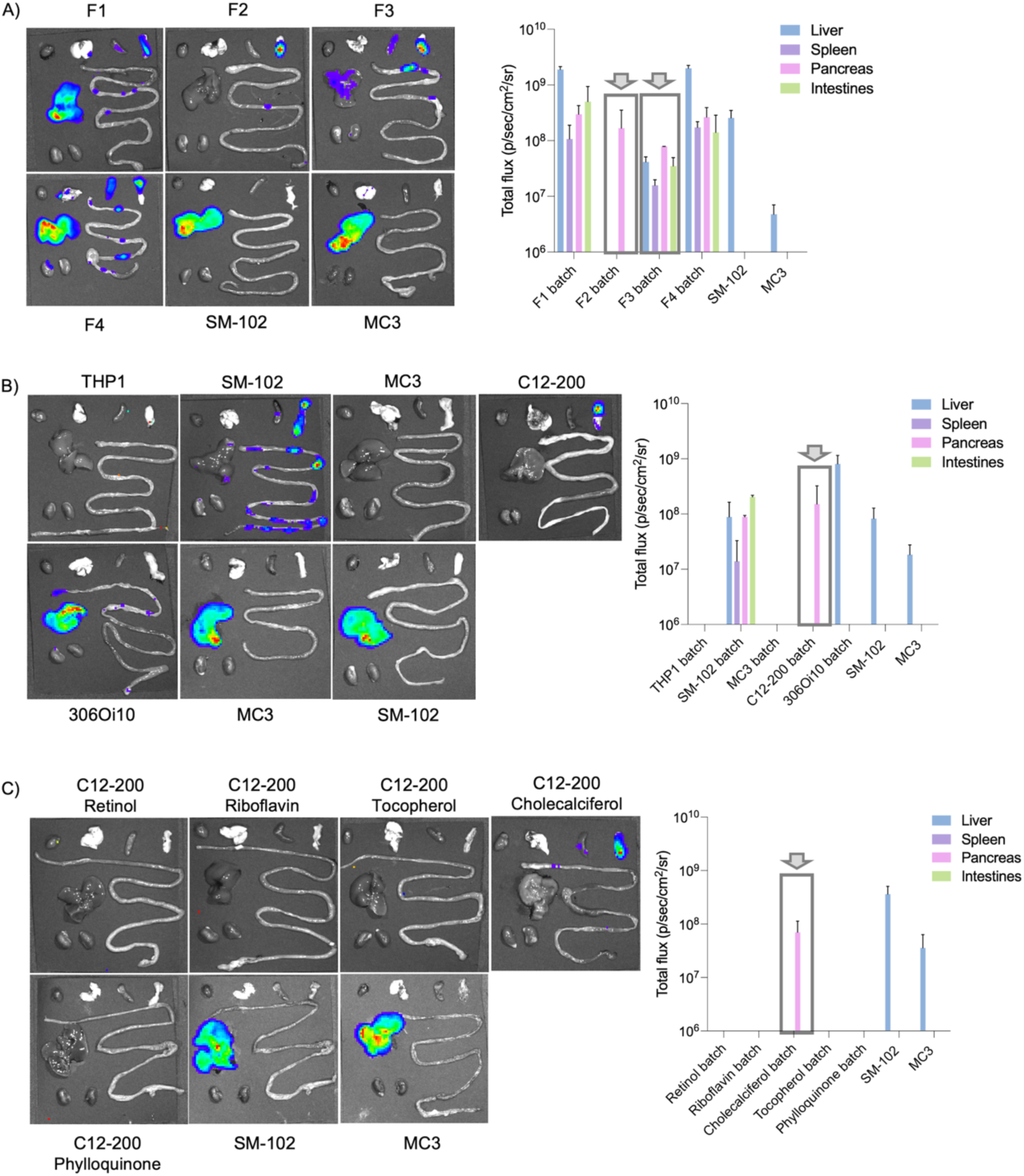
In Vivo screening of vitamins-based fifth component LNPs using batch analysis. **A)** IVIS images at 24h post-injection and graphical representation of total flux of FLuc mRNA fifth Component based LNPs. For the first screen, the LNPs were batched based on their formulation ratios (F1, F2, F3 and F4). SM-102 and MC3 were used as a control. C57BL/6 mice were injected intravenously with 0.5 mg/kg of pooled fifth Component-based LNPs. (n = 2 biologically independent mice, ± SD). **B)** IVIS images at 24h post-injection and graphical representation of total flux of FLuc mRNA fifth Component based LNPs. Pancreas targeting F2 and F3 LNPs were pooled together and batched based on the respective ionizable lipids (THP1, C12-200, SM-102, MC3, 306Oi10), and injected intravenously. SM-102 and MC3 were used as a control. C57BL/6 mice were injected intravenously with 0.5 mg/kg of pooled fifth Component-based LNPs LNPs. (n = 2 biologically independent mice, ± SD). **C)** IVIS images at 24h post-injection and graphical representation of total flux of FLuc mRNA fifth Component based LNPs. Selective pancreas targeting ionizable lipid C12-200 LNPs were then batched based on vitamins A (retinol), B2 (riboflavin), D3 (cholecalciferol), E (tocopherol), and K1 (phylloquinone). SM-102 and MC3 were used as a control. C57BL/6 mice were injected intravenously with 0.5 mg/kg of pooled fifth Component-based LNPs. (n = 2 biologically independent mice, ± SD).

To further validate our findings from the batch screening, we formulated C12-200 cholecalciferol F2 (C-CholF2) and C12-200 cholecalciferol F3 (C-CholF3) LNPs using a microfluidic device. Microfluidic mixing allows improved reproducibility by accurately mixing aqueous and ethanol phases, resulting in highly reproducible LNPs with desired physicochemical properties essential for clinical translation and large-scale production. The particle size and EE% of LNPs were further measured. C-CholF2 exhibited a particle size of 115.02 ± 1.66 nm, while the size of C-CholF3 was slightly lower (111.12 ± 1.45 nm). Moreover, the EE% was notably high, with 92.9% in the case of C-CholF2 and 96.8% in the case of C-CholF3, reflecting high mRNA encapsulation (**Figure 4A**). To validate the efficacy of these formulations in the pancreas, we intravenously injected each LNP at 0.5 mg/kg in C57BL/6 mice, along with the C12-200 control. The results showed that C-CholF3 had an average total flux of 1.04 × 10⁸ compared to 2.29 × 10⁷ for C-CholF2, indicating a 4.5-fold higher efficacy (**Figure 4B and S3)**. Thus, C-CholF3 exhibits a significantly enhanced ability to achieve targeted mRNA delivery to the pancreas, demonstrating a complete redirection of liver-targeting C12-200 LNPs with cholecalciferol (**Figure 4C**). To assess the dose-dependency of C-CholF3 and understand whether the protein expression in the pancreas could be regulated, we intravenously administered 0.25 mg/kg, 0.5 mg/kg, and 1 mg/kg of C-CholF3 LNPs to C57BL/6 mice and observed a distinguishable dose-dependent improvement in transfection efficiency **(Figure S4)**. This highlights the importance of optimizing dosing to maximize the efficiency of LNP-mediated mRNA delivery along with balancing safety at higher doses. These findings will be crucial in determining the therapeutic window and the dosing strategies of future clinical applications.

**Figure 4.**
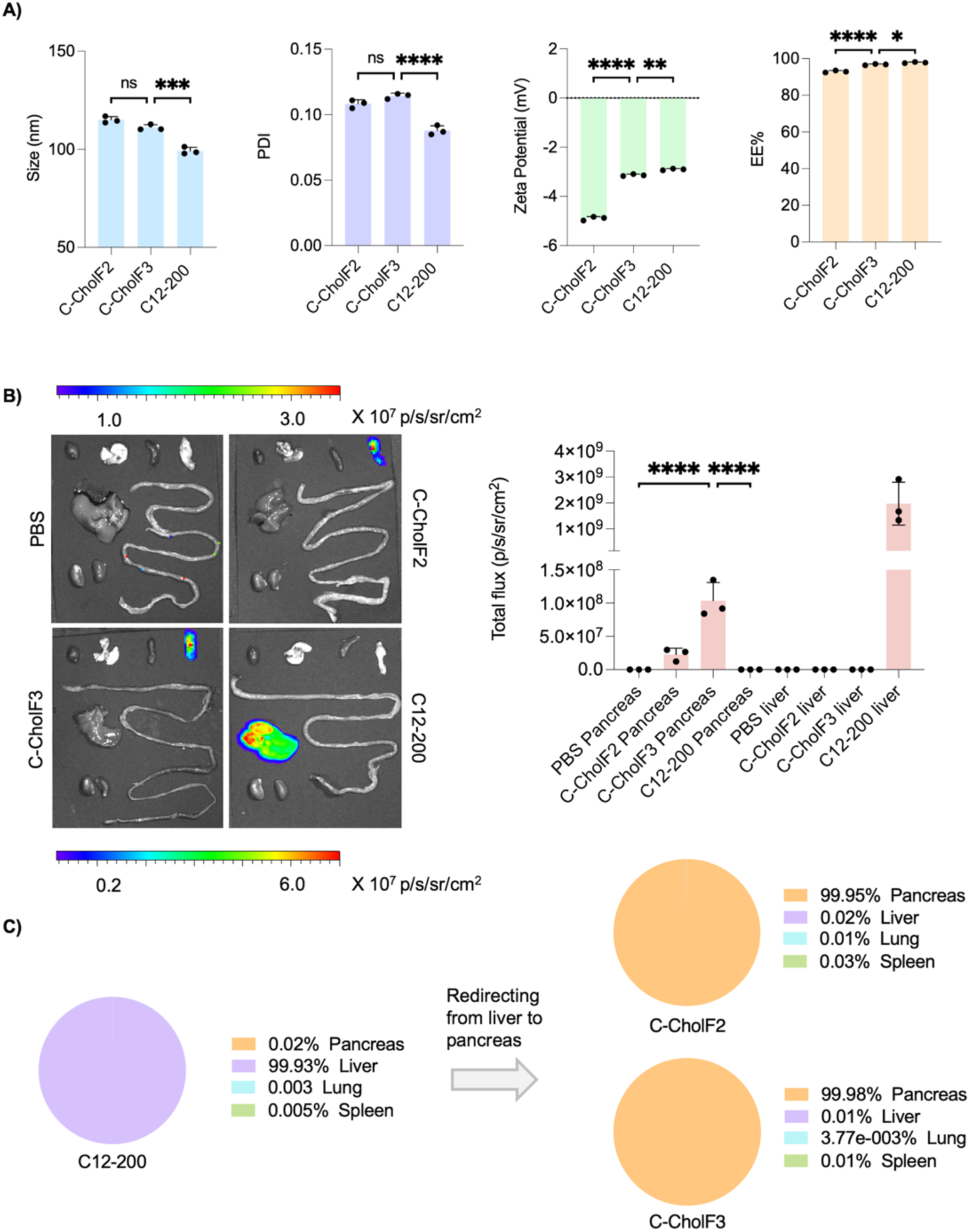
Validation of top vitamins-based fifth component LNPs, C-CholF2 and C-CholF3. **A)** Physiochemical properties such as size, PDI, ***ζ***-potential measurements, and mRNA encapsulation efficiency of C-CholF2 and C-CholF3 along with control (C12-200). (n = 3, **±** SD, **\***P < 0.05, **\*\***P < 0.01, **\*\*\***P < 0.001. NS, not significant, one-way ANOVA with Bonferroni posthoc analysis). **B)** Representative IVIS images at 24h post-injection and graphical representation of total flux of C-CholF2 and C-CholF3 mRNA LNPs injected intravenously at a dose of 0.5 mg/kg. PBS and C12-200 were also injected as a control (n=3 biologically independent mice, ± SD, *P < 0.05, **P < 0.01, ***P < 0.001. NS, not significant, one-way ANOVA with Bonferroni post-hoc analysis). **C)** Pie charts illustrating the percentage of protein expression occurring per organ after intravenous injection of C12-200 (four component traditional components) and direction to the pancreas for C-CholF2 and C-CholF3 mRNA LNPs (n=3 biologically independent mice).

Next, we investigated the in vivo biocompatibility of C-CholF3. H&E staining, both after 24 h and 48 h, showed no morphological damage or inflammatory response in pancreatic tissues treated with C-CholF3 compared to the controls treated with PBS (**Figure 5A, S5, and S6)**. The islet cells remained intact, and there was no evidence of necrosis or degeneration. Additionally, the liver tissues were examined with H&E staining after 48h and showed no significant change compared to the control. These findings highlight the safety and tolerability of C-CholF3 and its potential for future clinical applications. In addition to histopathological examination, the overall safety profile of C-CholF3 LNPs was studied by continuously monitoring body weight and hematological parameters. Over 21 days, no significant decrease in body weight was observed in mice **(Figure S7)**. The levels of liver enzymes such as ALT, AST, and alkaline phosphatase were within the normal range, with no renal toxicity observed based on the levels of BUN and CREA (**Figure 5B and S8)**. Proinflammatory cytokines, such as IL-1β, IL-6, and TNF-α, were measured 24 hours after injection and found to be comparable to PBS control, further confirming the low immunogenicity of the LNP (**Figure 5C**). Other immune biomarkers, including GM-CSF, IFNγ, IL-2, IL-4, IL-10, IL-12p70, and MCP-1, also showed comparable levels to control, indicating minimal systemic inflammatory response that is ideal for repeated dosing of LNPs **(Figure S9)**.

**Figure 5.**
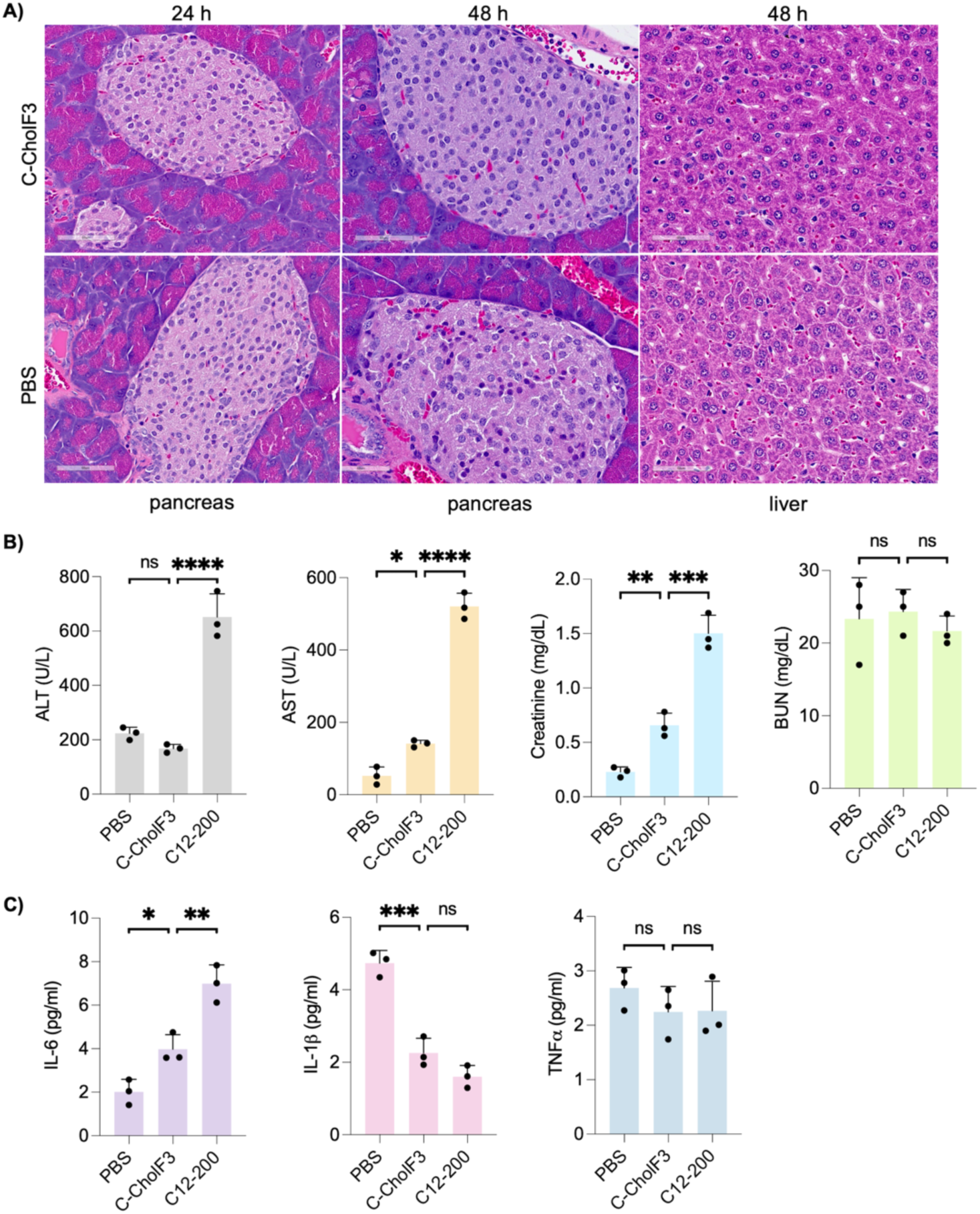
Toxicity and safety evaluation of C-CholF3 mRNA LNPs. **A)** Representative histology images of the pancreas (24h and 48h post-treatment) and liver sections 48h post-treatment with C-CholF3 encapsulating FLuc mRNA, and PBS (control) via intravenous injection in C57BL/6 mice at a dose of 0.5 mg/kg. Hematoxylin and Eosin (H&E) staining was performed, with images taken at 40× magnification. Scale bars: 60 μm (n = 3 biologically independent mice). **B)** Serum levels of liver enzymes, alanine aminotransferase (ALT) and aspartate aminotransferase (AST), renal parameters, blood urea nitrogen (BUN), and creatinine after intravenous administration with PBS, C-CholF3 and C12-200 mRNA LNPs at a dose of 0.5 mg/kg (n = 3, ± SD, *P < 0.05, **P < 0.01, ***P < 0.001. NS, not significant, one-way ANOVA with Bonferroni post-hoc analysis). **C)** Levels of IL-6, IL-1β, and TNFα in mice intravenously treated with C-CholF3 and C12-200 mRNA LNPs at a dose of 0.5 mg/kg (n=3 biologically independent mice, ± SD, *P< 0.05, **P< 0.01, ***P< 0.001; NS indicates no significance, one-way ANOVA with Bonferroni post-hoc analysis). PBS-injected mice were kept as the control group.

To ensure robustness and the stability of C-CholF3 mRNA LNPs for clinical applications, we stored them at 4°C and 20°C for various durations (1, 3, 7, and 21 days) and evaluated changes in their physiochemical properties and mRNA delivery efficacy. Throughout all storage conditions, the LNPs exhibited minimal changes in physiochemical properties, with parameters such as particle size, polydispersity index (PDI), ζ−potential, and EE% being consistent. Notably, the LNPs demonstrated robust FLuc expression even after 21 days of storage without any cryoprotectants, indicating exceptional stability **(Figure S10)**. This suggests that C-CholF3 LNPs can maintain their mRNA delivery potential over extended storage periods and can adopted for long-term storage and transport, making them suitable for clinical and commercial use. We also assessed the in vivo stability and kinetics of C-CholF3 mRNA LNPs to understand and determine their potential for extended release. For this, we administered the nanoparticles intravenously into C57BL/6 mice and monitored the bioluminescence signal over 72 hours. We observed that the bioluminescence intensity remained robust for up to 72 hours post-injection, indicating prolonged stability and expression of the delivered mRNA **(Figure S11)**. The higher stability of C-CholF3 LNPs could be advantageous for applications requiring prolonged gene expression in vivo.

Organ tropism has long been understood to be influenced by the apparent pKa of LNPs, with lower pKa values typically targeting the spleen and higher values favoring lung or liver delivery. However, pancreas-targeting LNPs had not been previously characterized. We evaluated the apparent pKa of C-CholF3 LNPs, which target the pancreas, and found it to be 7.31 compared to the C12-200 control at 6.71 **(Figure S12)**. This fine-tuning of pKa is crucial for the optimization of pancreas-specific LNP formulations. Overall, these findings provide a basis to conclude from our study that C-CholF3 LNPs represent a safe and highly effective modality of pancreas-targeted mRNA therapeutics with significant implications for pancreatic cancer and diabetes therapies. To further elucidate the mechanism of C-CholF3 mRNA delivery, we investigated the cellular internalization and subcellular trafficking of C-CholF3 LNPs by formulating fluorescently labeled particles using the lipid membrane anchor dye DiD and GFP mRNA. The labeled C-CholF3 LNPs were incubated with HMC3 cells at a dose of 125 ng per well in a 96-well plate to assess internalization efficiency and subcellular distribution over a 4-hour period. By the end of the incubation, the DiD signal, indicating the presence of the lipid nanoparticle, had largely diminished, while a strong GFP signal was detected in the majority of the cells. This transition in fluorescence from DiD to GFP indicates successful cellular uptake of the LNPs, followed by endosomal escape and efficient translation of the delivered mRNA (**Figure 6**). These findings suggest that C-CholF3 LNPs can promote rapid intracellular trafficking and successfully escape endosomal entrapment, a challenge that often limits the efficacy of LNP-based mRNA delivery systems. This characteristic may be advantageous for therapeutic applications that require quick and efficient protein expression.

**Figure 6.**
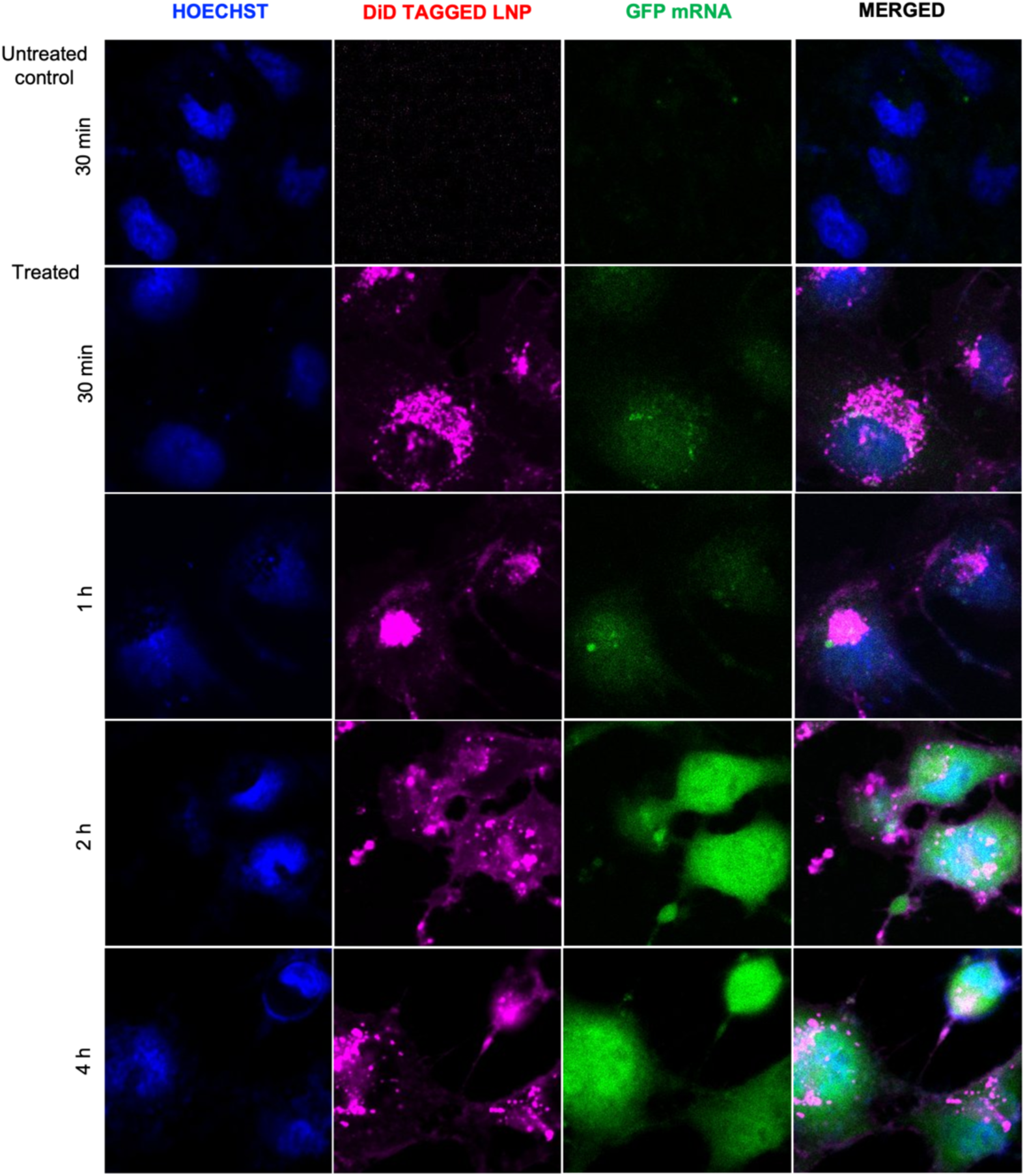
Cellular Uptake and Intracellular Trafficking of C-CholF3 LNPs. Confocal microscopy images showing the cellular uptake and intracellular trafficking of C-CholF3 lipid nanoparticles (LNPs) in HMC3 cells. The LNPs were labeled with DiD (magenta) to track their localization and encapsulated GFP mRNA for protein expression analysis. Time points of 30 minutes, 1 hour, 2 hours, and 4 hours post-treatment were evaluated to visualize nanoparticle dynamics and protein expression. Nuclei were stained with Hoechst 33342 (blue), and GFP protein expression (green) was observed as an indicator of successful mRNA translation. Scale bars represent 20 μm.

Having demonstrated that C-CholF3 can deliver mRNA to the pancreas, we next utilized the Ai14 mouse model to assess the tissue-specific gene-editing capability of C-CholF3 LNPs. The Ai14 mice are genetically engineered and contain a construct designed for Cre-mediated gene editing, featuring a LoxP-stop-LoxP cassette downstream of the CAG promoter. After successful delivery of Cre recombinase, the stop cassette is excised, activating tdTomato fluorescence in the targeted cells.^[57,58]^ Thus, the system simplifies the identification and quantification of the gene-edited cells, as the cells with Cre mRNA expression exhibit characteristic red tdTomato fluorescence in a tissue-specific manner (**Figure 7A**). We formulated the C-CholF3 LNPs with Cre recombinase mRNA and intravenously injected them into the Ai14 mice at 1.5 mg/kg. After 120 h, we imaged fluorescence in various tissues using the IVIS and observed a very strong fluorescent signal in the pancreas (**Figure 7B**) with more than 99% selectivity toward pancreas (**Figure 7C and S13)**. All other tissues generally show very minimal fluorescence, suggesting that this formulation limits gene editing mostly to the pancreas-further reinforcing this as a targeted delivery vehicle.

**Figure 7.**
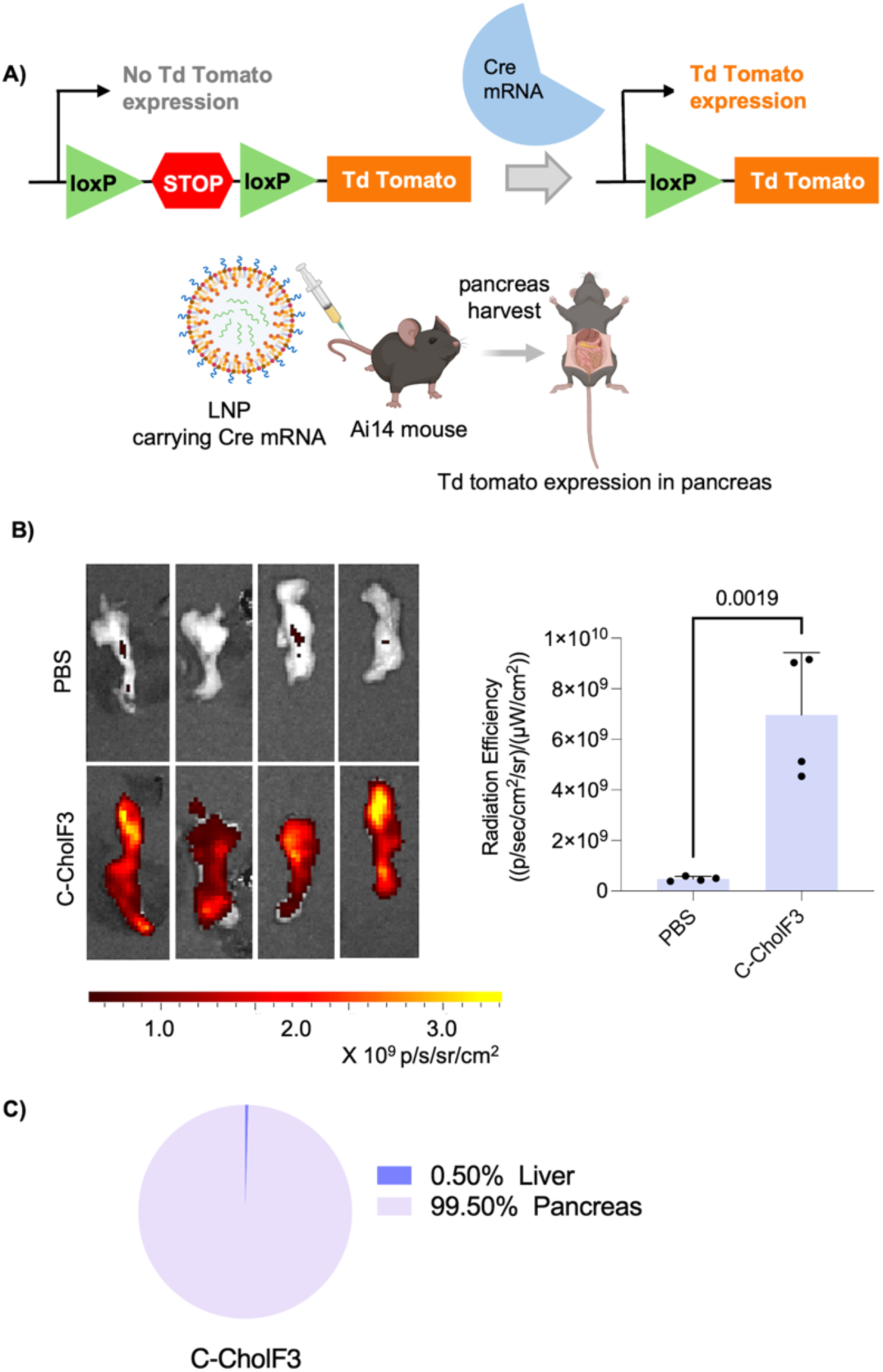
Efficient and tissue-specific tdTomato expression in the pancreas with C-CholF3 LNPs. **A)** Schematic illustration of Cre mRNA delivery and subsequent Cre-mediated removal of the stop cassette, leading to the activation of tdTomato expression in the Ai14 Cre-loxP mouse model. LNPs formulated with Cre mRNA were injected intravenously and the pancreas was imaged using the IVIS. **B)** Representative tdTomato expression in the pancreas 120h post-injection and graphical representation of total flux of C-CholF3 LNPs containing Cre mRNA, and PBS treated control injected intravenously to Ai14 mice at a dose of 1.5 mg/kg (n=4 biologically independent mice, ± SD, *P < 0.05, **P < 0.01, ***P < 0.001. NS, not significant, one-way ANOVA with Bonferroni post-hoc analysis). **C)** Pie charts illustrating the percentage of protein expression occurring in pancreas and liver after intravenous injection of Cre mRNA containing C-CholF3 mRNA LNPs. (n=3 biologically independent mice).

## Conclusion

In summary, our study presents a comprehensive investigation of C-CholF3 LNPs as a novel mRNA delivery platform. We developed a diverse library of 100 LNPs, incorporating various vitamins as a fifth component, and evaluated their transfection efficacy both in vitro and in vivo. Through systematic batch analysis, we identified two key formulations, C-CholF2 and C-CholF3, which contained cholecalciferol as a fifth component. Remarkably, these formulations demonstrated selective (> 99%) and robust protein expression in the pancreas with significantly lower toxicity than parent 4 component C12-200. Furthermore, C-CholF3 also demonstrated robust pancreas-specific TdTomato expression in the Ai14 transgenic mouse model, highlighting the effectiveness of C-CholF3 LNPs in gene editing. This indicates that formulations with the right endogenous fifth component could be utilized to redirect therapies to extrahepatic space with minimal toxicity, potentially advancing the development of gene therapy or personalized medicine for pancreatic diseases with the capability of repeat administration.

## Experimental section

### Animal experiments

All animal procedures were conducted in accordance with guidelines and approval from the Institutional Animal Care and Use Committee (IACUC) at the University of Las Vegas, Nevada (protocol #01218). Female and male C57BL/6J mice (6–8 weeks old, approximately 20 g) and B6.Cg-Gt(ROSA)26Sortm14(CAG-tdTomato)Hze/J mice (6–8 weeks old, approximately 20 g) were obtained from Jackson Laboratory (Bar Harbor, ME, USA). Mice were injected with LNPs formulated with FLuc mRNA at a dose of 0.5 mg/kg via the lateral tail vein. A D-luciferin solution (30 mg/mL in 1X PBS; PerkinElmer) was prepared, and mice were injected intraperitoneally with 130 µL of this solution after 24 hours (for intravenous injections). After a 10-minute incubation, the mice were euthanized using CO2, and organs (liver, spleen, kidney, pancreas, heart, lung) were harvested and imaged with an in vivo imaging system (IVIS; PerkinElmer, Waltham, MA, USA). Luminescence flux was quantified using Living Image Software (PerkinElmer). For toxicity assessment, blood samples were collected via cardiac puncture 24 hours after treatment. Mice were then sacrificed by CO2 asphyxiation, and organs were collected. Blood samples were allowed to coagulate for 20 minutes at room temperature and centrifuged at 2000 x g for 20 minutes at 4°C to obtain high-quality of serum. Serum liver enzyme levels were measured by VRL Animal Health Diagnostics and cytokine panel was performed by Eve Technologies Corp. (Calgary, Alberta). For Ai14 mice experiments, fluorescence was quantified at an excitation/emission of 554/581 nm. Regions of interest (ROIs) of a constant size were placed around each organ’s image, and total luminescence flux and radiant efficiency were reported as mean ± standard deviation.

Statistics: Statistical analysis of the results was performed by One-Way ANOVA followed by Bonferroni post-hoc analysis to compare multiple replicate means using Prism 10 (GraphPad). Differences were considered significant when p < 0.05.

## Supporting Information

Supporting Information is available from the Wiley Online Library or from the author.

## Acknowledgments

This work was supported by the College of Sciences at the University of Nevada Las Vegas and the PhRMA Foundation Research Starter Grant in Drug Delivery 2024 FSGDL 1170646. We acknowledge the use of resources at the Nevada Institute of Personalized Medicine (NIPM) Core Facility. S.P. acknowledges support from the National Institute on Aging (NIA) award number K25AG070286. The TOC and **Figures 1A, 1B**, and **6A** were partly created using BioRender.com.

## Author Contributions

I.I. and C.B. conceived the project, designed the experiments, and wrote the manuscript. I.I. synthesized the LNPs, performed all the in vitro and in vivo experiments, and analyzed data. L.P. assisted I.I. with the high-throughput screen and N.T. with immunofluorescence optimization. A.S. helped with confocal imaging. P.G. and S.P. provided valuable comments. C.B. directed the research.

## References

[1] T. Vavilis, E. Stamoula, A. Ainatzoglou, A. Sachinidis, M. Lamprinou, I. Dardalas, I. S. Vizirianakis, Pharmaceutics 2023, 15, 166.

[2] R. Gambaro, I. Rivero Berti, M. J. Limeres, C. Huck-Iriart, M. Svensson, S. Fraude, L. Pretsch, S. Si, I. Lieberwirth, S. Gehring, M. Cacicedo, G. A. Islan, Pharmaceutics 2024, 16, 771.

[3] R. J. Chandler, Proceedings of the National Academy of Sciences 2019, 116, 20804.

[4] R. Verbeke, I. Lentacker, S. C. De Smedt, H. Dewitte, Journal of Controlled Release 2021, 333, 511.

[5] L. Zhang, K. R. More, A. Ojha, C. B. Jackson, B. D. Quinlan, H. Li, W. He, M. Farzan, N. Pardi, H. Choe, npj Vaccines 2023, 8, 1.

[6] E. H. Pilkington, E. J. A. Suys, N. L. Trevaskis, A. K. Wheatley, D. Zukancic, A. Algarni, H. Al-Wassiti, T. P. Davis, C. W. Pouton, S. J. Kent, N. P. Truong, Acta Biomaterialia 2021, 131, 16.

[7] L. Schoenmaker, D. Witzigmann, J. A. Kulkarni, R. Verbeke, G. Kersten, W. Jiskoot, D. J. A. Crommelin, International Journal of Pharmaceutics 2021, 601, 120586.

[8] E. Álvarez-Benedicto, L. Farbiak, M. M. Ramírez, X. Wang, L. T. Johnson, O. Mian, E. D. Guerrero, D. J. Siegwart, Biomaterials Science 2022, 10, 549.

[9] C. Hald Albertsen, J. A. Kulkarni, D. Witzigmann, M. Lind, K. Petersson, J. B. Simonsen, Adv Drug Deliv Rev 2022, 188, 114416.

[10] R. Zhang, R. El-Mayta, T. J. Murdoch, C. C. Warzecha, M. M. Billingsley, S. J. Shepherd, N. Gong, L. Wang, J. M. Wilson, D. Lee, M. J. Mitchell, Biomaterials Science 2021, 9, 1449.

[11] M. Kim, M. Jeong, S. Hur, Y. Cho, J. Park, H. Jung, Y. Seo, H. A. Woo, K. T. Nam, K. Lee, H. Lee, Science Advances 2021, 7, eabf4398.

[12] M. M. Żak, L. Zangi, Pharmaceutics 2021, 13, 1675.

[13] R. Pattipeiluhu, G. Arias-Alpizar, G. Basha, K. Y. T. Chan, J. Bussmann, T. H. Sharp, M.-A. Moradi, N. Sommerdijk, E. N. Harris, P. R. Cullis, A. Kros, D. Witzigmann, F. Campbell, Advanced Materials 2022, 34, 2201095.

[14] T. Wei, Y. Sun, Q. Cheng, S. Chatterjee, Z. Traylor, L. T. Johnson, M. L. Coquelin, J. Wang, M. J. Torres, X. Lian, X. Wang, Y. Xiao, C. A. Hodges, D. J. Siegwart, Nat Commun 2023, 14, 7322.

[15] Q. Cheng, T. Wei, L. Farbiak, L. T. Johnson, S. A. Dilliard, D. J. Siegwart, Nat. Nanotechnol. 2020, 15, 313.

[16] X. Wang, S. Liu, Y. Sun, X. Yu, S. M. Lee, Q. Cheng, T. Wei, J. Gong, J. Robinson, D. Zhang, X. Lian, P. Basak, D. J. Siegwart, Nat Protoc 2023, 18, 265.

[17] D. Chen, N. Parayath, S. Ganesh, W. Wang, M. Amiji, Nanoscale 2019, 11, 18806.

[18] X. Jiao, X. He, S. Qin, X. Yin, T. Song, X. Duan, H. Shi, S. Jiang, Y. Zhang, X. Song, WIREs Nanomedicine and Nanobiotechnology 2024, 16, e1992.

[19] X. Han, M.-G. Alameh, K. Butowska, J. J. Knox, K. Lundgreen, M. Ghattas, N. Gong, L. Xue, Y. Xu, M. Lavertu, P. Bates, J. Xu, G. Nie, Y. Zhong, D. Weissman, M. J. Mitchell, Nat. Nanotechnol. 2023, 18, 1105.

[20] J. Qin, L. Xue, N. Gong, H. Zhang, S. J. Shepherd, R. M. Haley, K. L. Swingle, M. J. Mitchell, RSC Advances 2022, 12, 25397.

[21] M. Herrera-Barrera, R. C. Ryals, M. Gautam, A. Jozic, M. Landry, T. Korzun, M. Gupta, C. Acosta, J. Stoddard, R. Reynaga, W. Tschetter, N. Jacomino, O. Taratula, C. Sun, A. K. Lauer, M. Neuringer, G. Sahay, Science Advances 2023, 9, eadd4623.

[22] B. Elibol-Can, N. Simsek-Ozek, F. Severcan, M. Severcan, E. Jakubowska-Dogru, British Journal of Nutrition 2015, 113, 45.

[23] S. S. Khairnar, K. R. Surana, E. D. Ahire, S. K. Mahajan, D. M. Patil, D. D. Sonawane, in Vitamins as Nutraceuticals, John Wiley & Sons, Ltd, 2023, pp. 35–60.

[24] P. Singh, R. K. Kesharwani, R. K. Keservani, in Sustained Energy for Enhanced Human Functions and Activity (Ed.: D. Bagchi), Academic Press, 2017, pp. 385–407.

[25] M. S. Aurora-Prado, C. A. Silva, M. F. M. Tavares, K. D. Altria, Chroma 2010, 72, 687.

[26] P. Anvith, R. Sankar, 2015, 12.

[27] S. L. Stevens, Nursing Clinics 2021, 56, 33.

[28] S. H. Kiaie, N. Majidi Zolbanin, A. Ahmadi, R. Bagherifar, H. Valizadeh, F. Kashanchi, R. Jafari, J Nanobiotechnol 2022, 20, 276.

[29] Y. Zong, Y. Lin, T. Wei, Q. Cheng, Advanced Materials 2023, 35, 2303261.

[30] H. Tanaka, R. Miyama, Y. Sakurai, S. Tamagawa, Y. Nakai, K. Tange, H. Yoshioka, H. Akita, Pharmaceutics 2021, 13, 2097.

[31] S. Wu, L. Lin, L. Shi, S. Liu, WIREs Nanomedicine and Nanobiotechnology 2024, 16, e1978.

[32] S. Roy, A. Awasthi, in Nutrition and Immunity (Eds.: M. Mahmoudi, N. Rezaei), Springer International Publishing, Cham, 2019, pp. 53–73.

[33] M. R. Bono, G. Tejon, F. Flores-Santibañez, D. Fernandez, M. Rosemblatt, D. Sauma, Nutrients 2016, 8, 349.

[34] D. Julian McClements, Food & Function 2018, 9, 22.

[35] S. Balasubramaniam, J. Yaplito-Lee, jtgg 2020, 4, 285.

[36] J. T. Pinto, J. Zempleni, Advances in Nutrition 2016, 7, 973.

[37] K. Wang, M. Dong, W. Sheng, Q. Liu, D. Yu, Q. Dong, Q. Li, J. Wang, Histopathology 2015, 67, 386.

[38] M. Infante, C. Ricordi, J. Sanchez, M. J. Clare-Salzler, N. Padilla, V. Fuenmayor, C. Chavez, A. Alvarez, D. Baidal, R. Alejandro, M. Caprio, A. Fabbri, Nutrients 2019, 11, 2185.

[39] U. Singh, S. Devaraj, I. Jialal, Annual Review of Nutrition 2005, 25, 151.

[40] C. Bolton-Smith, M. E. McMurdo, C. R. Paterson, P. A. Mole, J. M. Harvey, S. T. Fenton, C. J. Prynne, G. D. Mishra, M. J. Shearer, Journal of Bone and Mineral Research 2007, 22, 509.

[41] M. Halder, P. Petsophonsakul, A. C. Akbulut, A. Pavlic, F. Bohan, E. Anderson, K. Maresz, R. Kramann, L. Schurgers, International Journal of Molecular Sciences 2019, 20, 896.

[42] C. J. Calvente, A. Sehgal, M. Diken, D. Schuppan, Hepatology 2013, 58, 583A.

[43] C. Huang, Y. Zhang, J. Su, X. Guan, S. Chen, X. Xu, X. Deng, L. Zhang, J. Huang, International Journal of Nanomedicine 2023, 18, 7785.

[44] F. Ferraresso, A. W. Strilchuk, L. J. Juang, L. G. Poole, J. P. Luyendyk, C. J. Kastrup, Mol. Pharmaceutics 2022, 19, 2175.

[45] I. Isaac, A. Shaikh, M. Bhatia, Q. Liu, S. Park, C. Bhattacharya, ACS Nano 2024, 18, 29045.

[46] S. Qin, X. Tang, Y. Chen, K. Chen, N. Fan, W. Xiao, Q. Zheng, G. Li, Y. Teng, M. Wu, X. Song, Sig Transduct Target Ther 2022, 7, 1.

[47] B. J. Henriques, T. G. Lucas, C. M. Gomes, Current Drug Targets 2016, 17, 1527.

[48] G. Xu, M. Redard, G. Gabbiani, P. Neuville, The American Journal of Pathology 1997, 151, 1741.

[49] R. H. Böger, S. M. Bode-Böger, L. Phivthong-ngam, R. P. Brandes, E. Schwedhelm, A. Mügge, M. Böhme, D. Tsikas, J. C. Frölich, Atherosclerosis 1998, 141, 31.

[50] D. A. Faherty, A. Bendich, in Methods in Enzymology, Academic Press, 1990, pp. 252–259.

[51] H.-M. Lo, S.-W. Wang, C.-L. Chen, P.-H. Wu, W.-B. Wu, Food & Function 2014, 5, 140.

[52] I. K. Midha, N. Kumar, A. Kumar, T. Madan, Reviews in Medical Virology 2021, 31, e2204.

[53] G. Scalabrino, Journal of Neurochemistry 2009, 111, 1309.

[54] K. Dakshinamurti, Vitamin Receptors: Vitamins as Ligands in Cell Communication, Cambridge University Press, 1994.

[55] Y. Shaik-Dasthagirisaheb, G. Varvara, G. Murmura, A. Saggini, A. Caraffa, P. Antinolfi, S. Tete, D. Tripodi, F. Conti, E. Cianchetti, E. Toniato, M. Rosati, L. Speranza, A. Pantalone, R. Saggini, M. Tei, A. Speziali, P. Conti, T. Theoharides, F. Pandolfi, Journal of biological regulators and homeostatic agents 2012, 27, 291.

[56] S. Aslam, 2020.

[57] L. Xue, A. G. Hamilton, G. Zhao, Z. Xiao, R. El-Mayta, X. Han, N. Gong, X. Xiong, J. Xu, C. G. Figueroa-Espada, S. J. Shepherd, A. J. Mukalel, M.-G. Alameh, J. Cui, K. Wang, A. E. Vaughan, D. Weissman, M. J. Mitchell, Nat Commun 2024, 15, 1884.

[58] J. Tuma, Y.-J. Chen, M. G. Collins, A. Paul, J. Li, H. Han, R. Sharma, N. Murthy, H. Y. Lee, Biochemistry 2023, 62, 3533.

