## Supplementary material for "Reengineering mRNA lipid nanoparticles for systemic delivery to pancreas": Supplementary Information.docx

|  | **Ionizable lipid** | **DOPE** | **Cholesterol** | **PEG** | **5th Component** |
| --- | --- | --- | --- | --- | --- |
| **F1** | 30 | 16 | 46.5 | 2.5 | 5 |
| **F2** | 25 | 16 | 46.5 | 2.5 | 10 |
| **F3** | 20 | 16 | 46.5 | 2.5 | 15 |
| **F4** | 35 | 16 | 41.5 | 2.5 | 5 |

**Table S1.** Formulation ratios used to synthesize fifth component-based mRNA LNPs

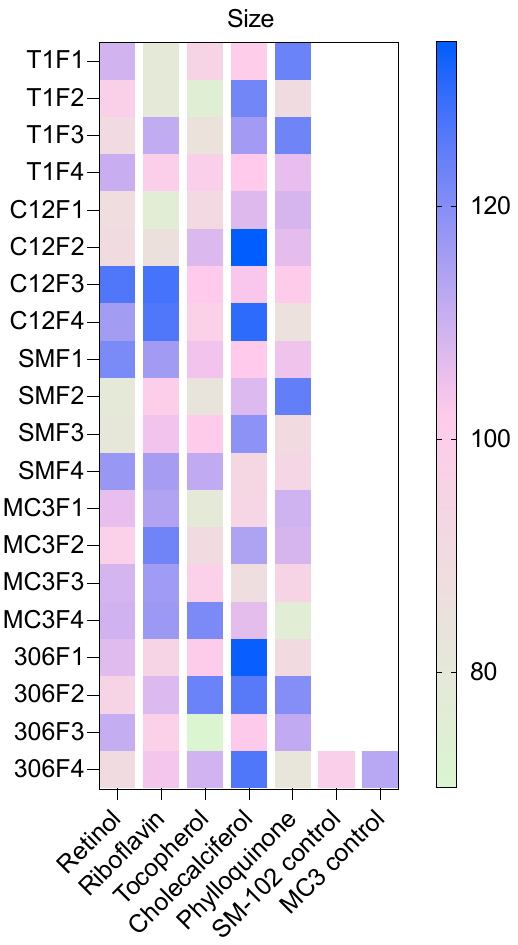

**Figure S1.** Heatmap representing size (in nm) of the fifth component-based mRNA LNPs measured using DLS (n=3).

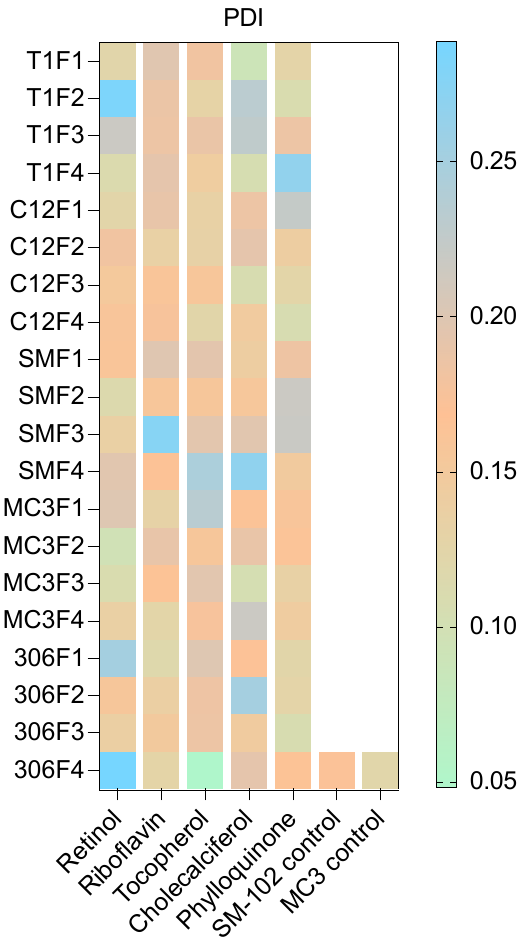

**Figure S2.** Heatmap representing PDI of the fifth-component mRNA LNPs measured using DLS (n=3).

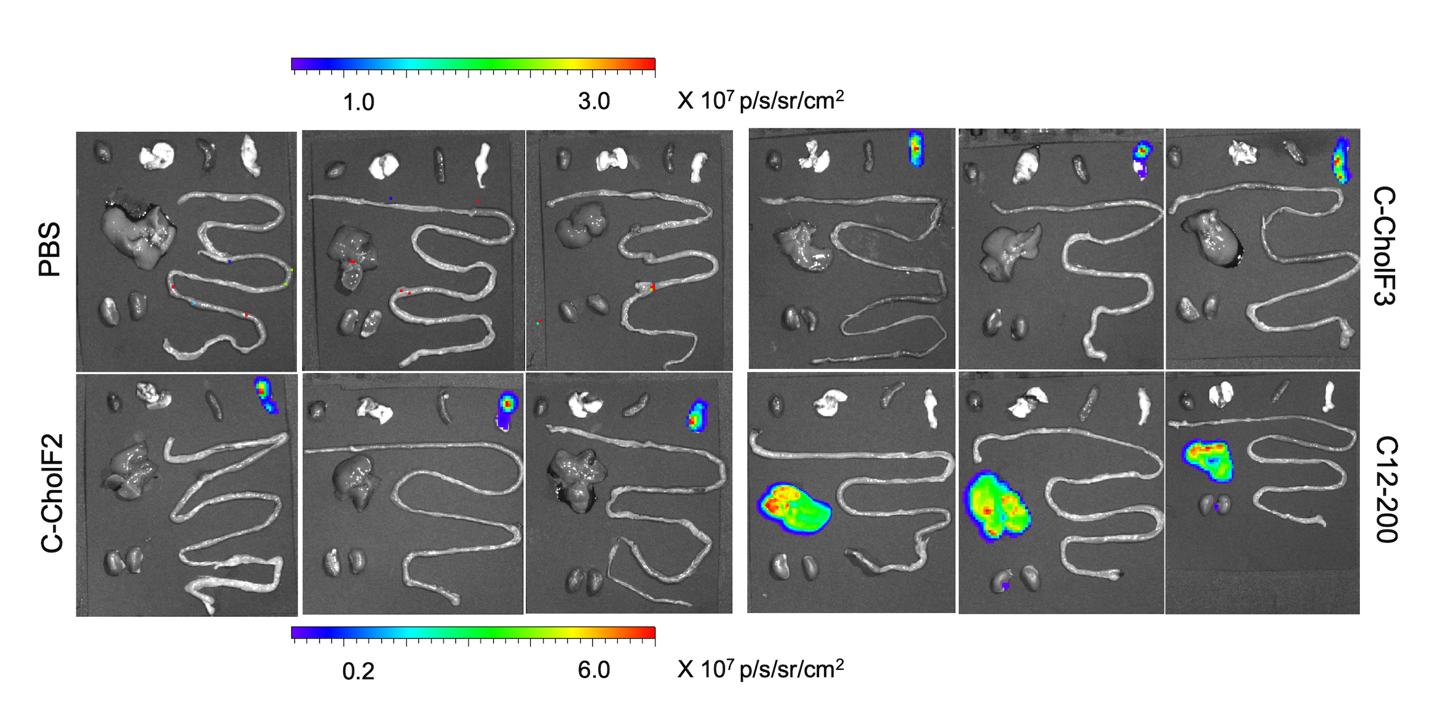

**Figure S3.** All replicates of IVIS images from **Figure 4B** after administration of C-CholF2 and C-CholF3 mRNA LNPs injected intravenously at a dose of 0.5 mg/kg. PBS and C12-200 were also injected (n=3 biologically independent mice).

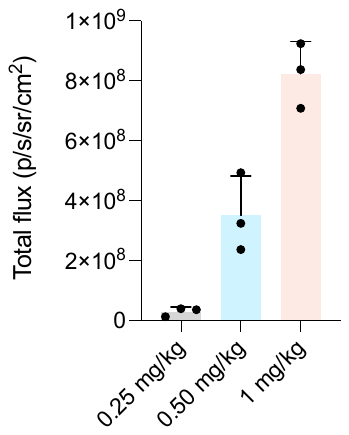

**Figure S4.** Graphical representation of total flux at 24h post-injection of different doses (0.25, 0.5, and 1 mg/kg) of C-CholF3  mRNA LNPs administered intravenously in C57BL/6 mice (n = 3 biologically independent mice).

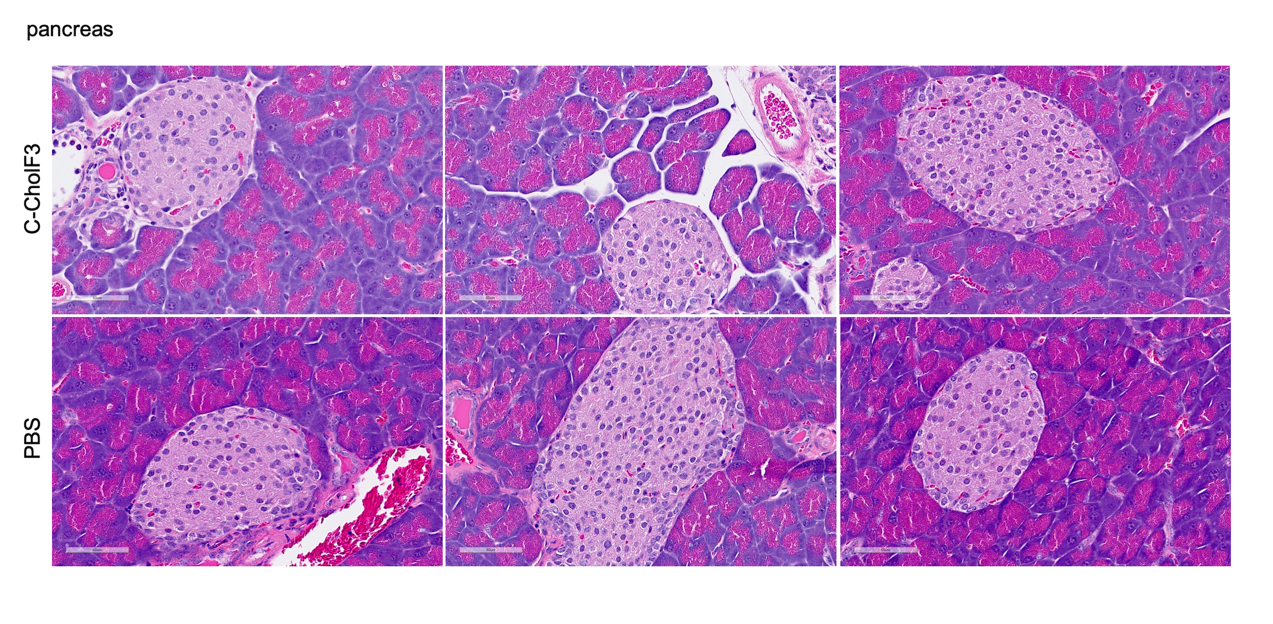

**Figure S5.** Replicates of H&E staining of the pancreas after 24 h used to generate **Figure 5A.**

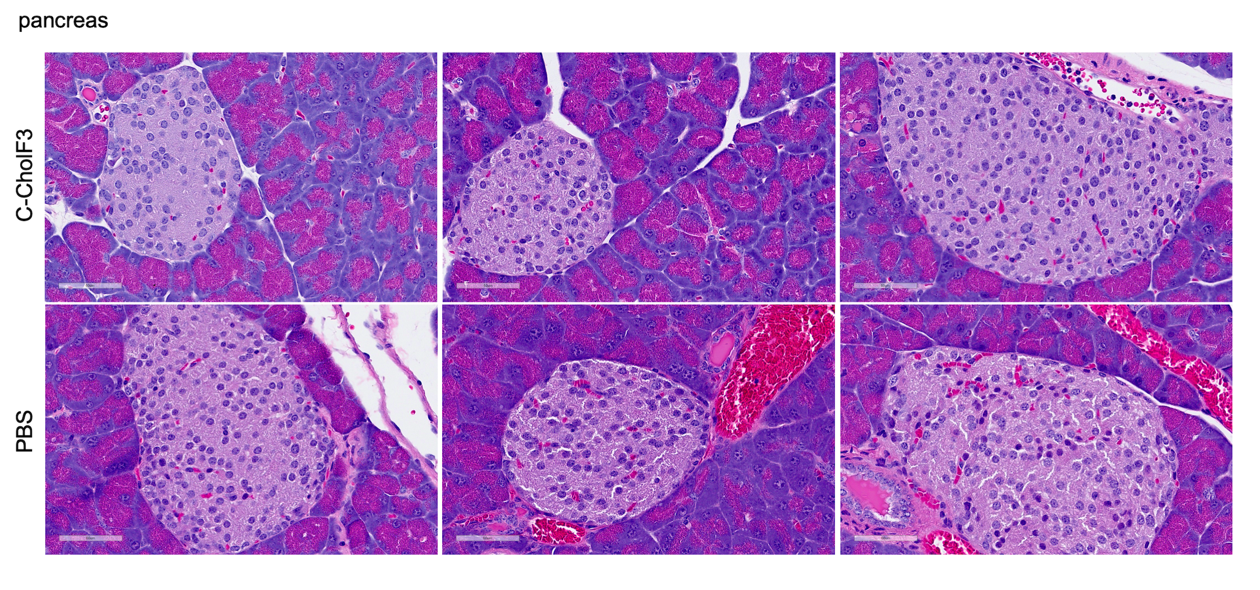

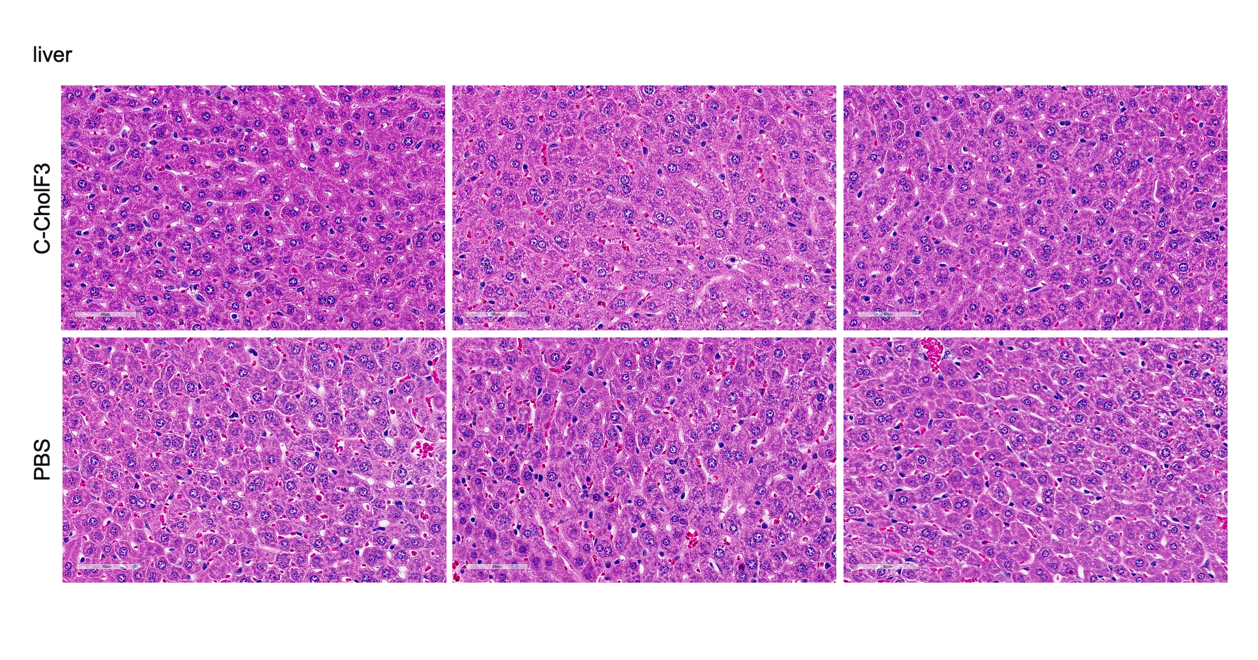

**Figure S6.** Replicates of H&E staining of the pancreas and liver after 24 h used to generate **Figure 5A.**

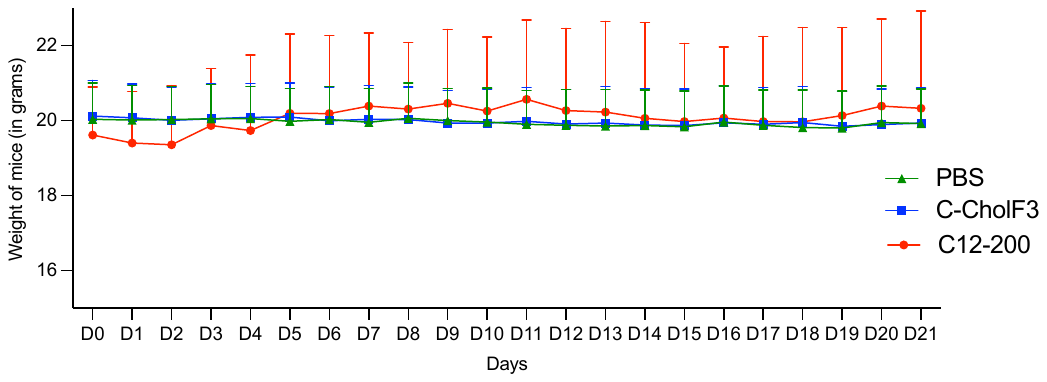

**Figure S7.** The body weight of mice treated with C-CholF3 and C12-200 LNPs at a dose of 0.5 mg/kg (i.v.) for 21 days (n=3). PBS was intravenously injected into mice as a control.

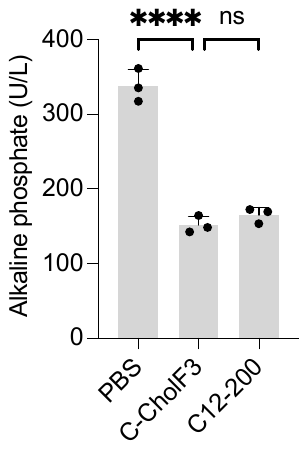

**Figure S8.** Serum levels of the liver enzyme alkaline phosphatase, after intravenous administration with PBS, C-CholF3, and C12-200 mRNA LNPs at a dose of 0.5 mg/kg. (n=3 biologically independent mice, ± SD, *P < 0.05, **P < 0.01, ***P < 0.001. NS, not significant, one-way ANOVA with Bonferroni post-hoc analysis).

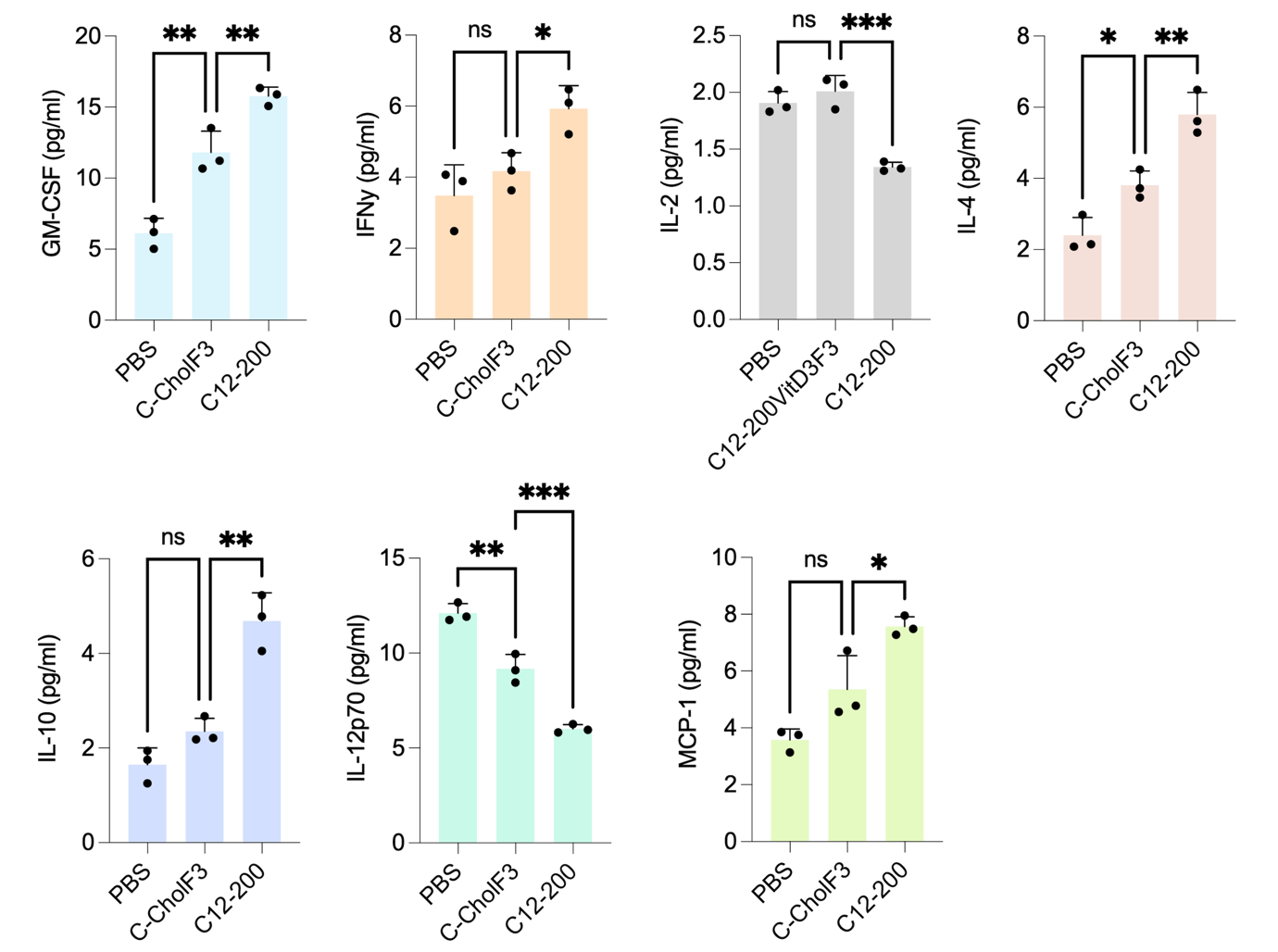

**Figure S9.** Levels of biomarkers such as GM-CSF, IFNγ, IL-2, IL-4, IL-10, IL-12P70, and MCP-1  in mice intravenously treated with C-CholF3 and C12-200 LNPs at a dose of 0.5 mg/kg (n=3 biologically independent mice, ± SD, *P < 0.05, **P < 0.01, ***P < 0.001. NS, not significant, one-way ANOVA with Bonferroni post-hoc analysis). PBS-injected mice were kept as the control group.

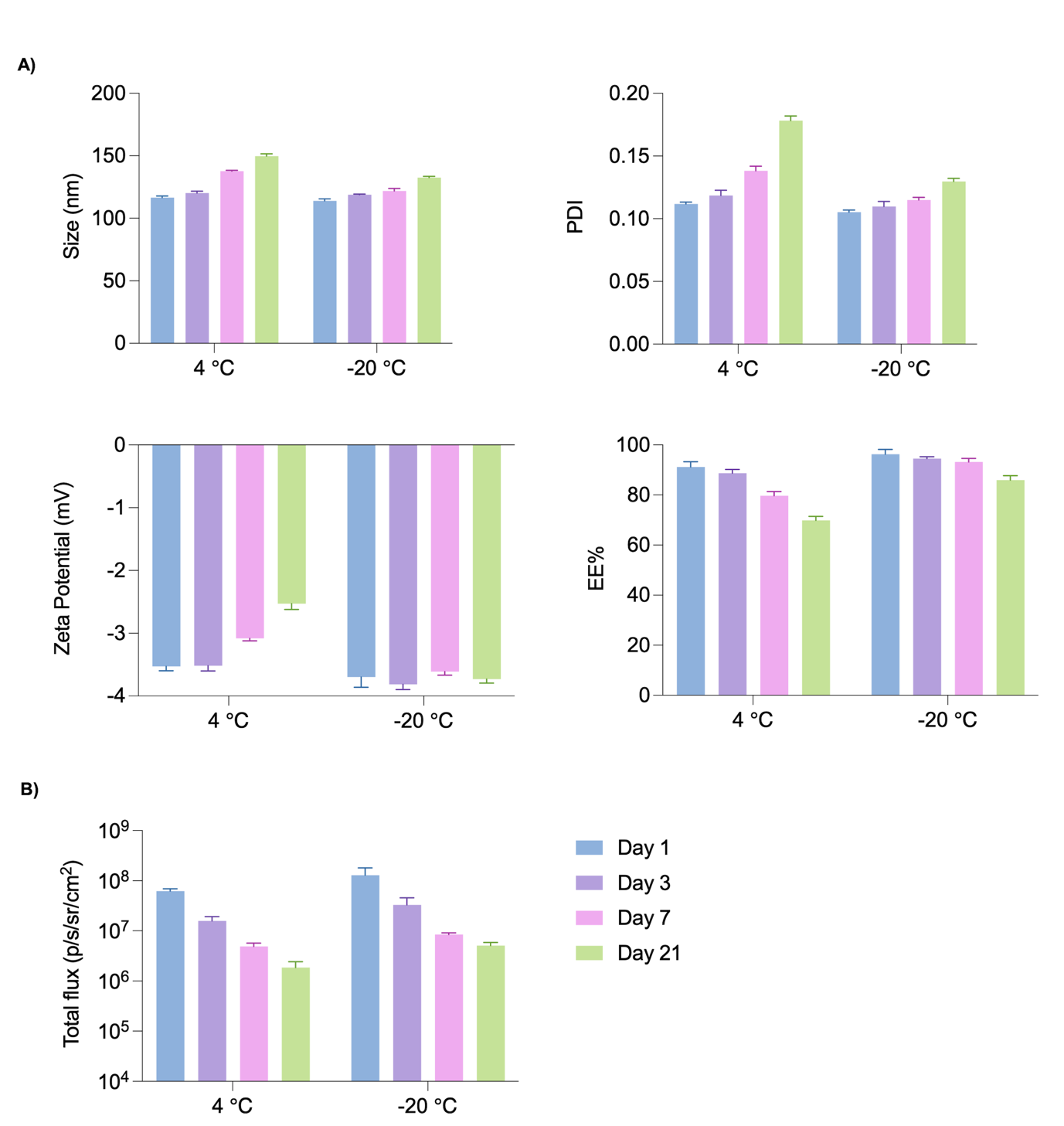

**Figure S10.** Stability study of C-CholF3 mRNA LNPs. A) Size, PDI, and ζ-potential of C-CholF3 LNPs under different storage conditions (4°C and -20°C) for 1, 3, 7, and 21 days. B) Representative IVIS images of total flux 24 h after intravenous injection of FLuc mRNA-loaded C-CholF3  LNPs after being stored at different temperatures for extended time (0.5 mg/kg, n = 3).

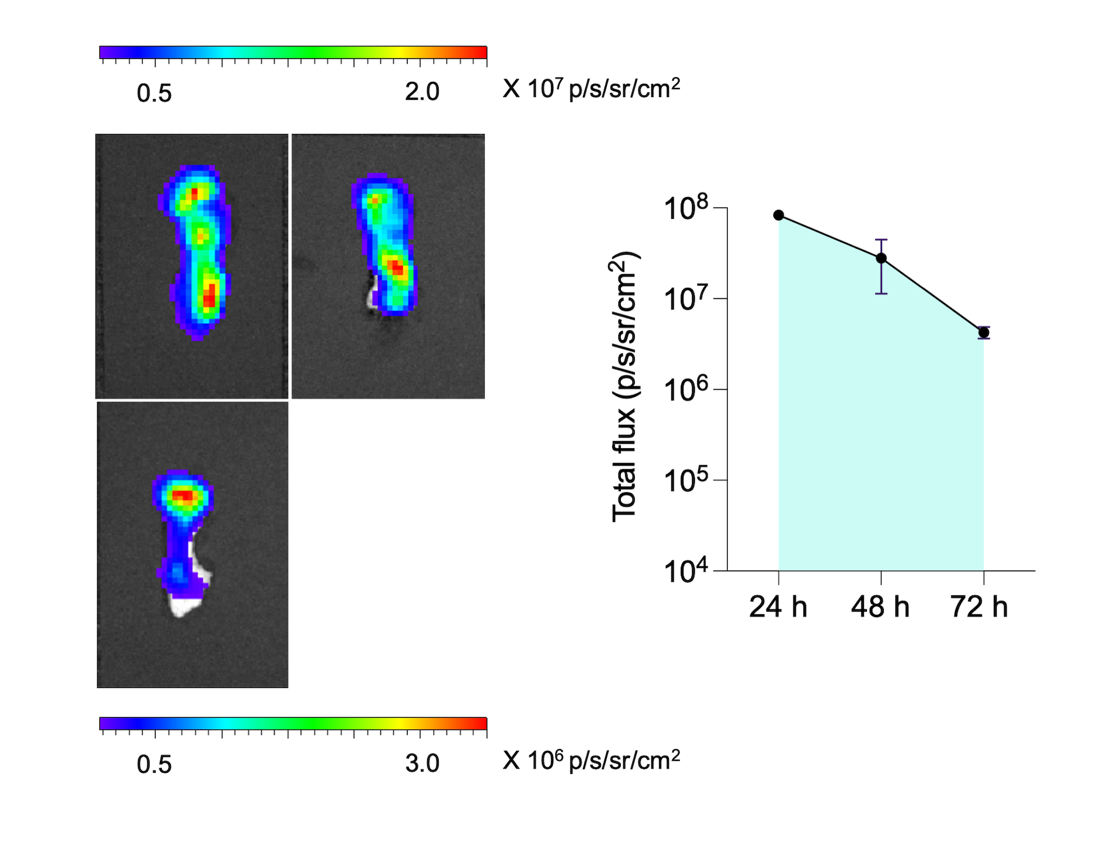

**Figure S11.** IVIS images and graphical representation of in vivo kinetics of FLuc expression following intravenous injection of C-CholF3 mRNA LNPs at a dose of 0.5 mg/kg (n = 3 biologically independent mice, ± SD). The luciferase expression was visualized at 24, 48, and 72 hours after injection by IVIS.

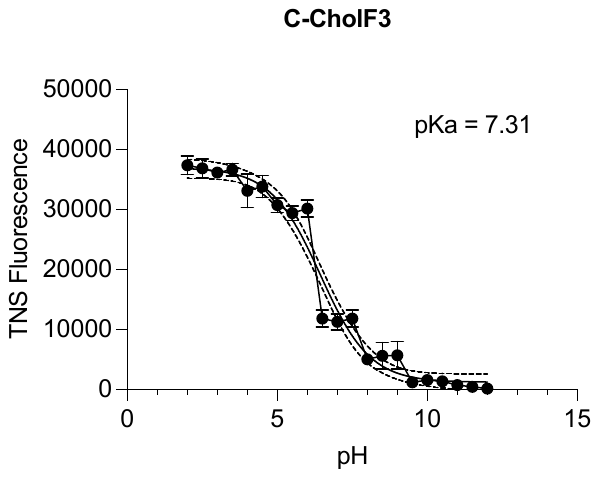

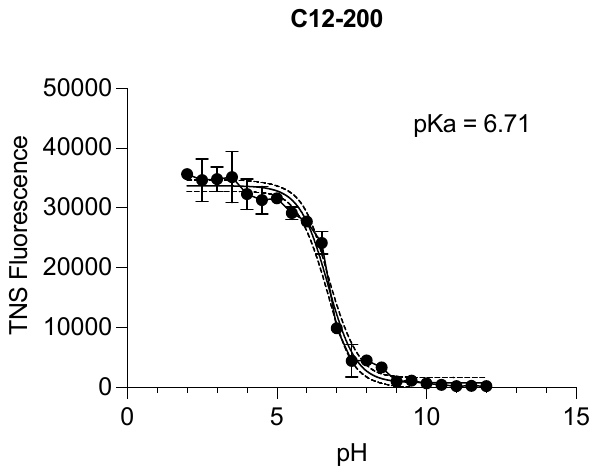

**Figure S12.** TNS curves for pKa measurements of C-CholF3 and C12-200 LNPs.

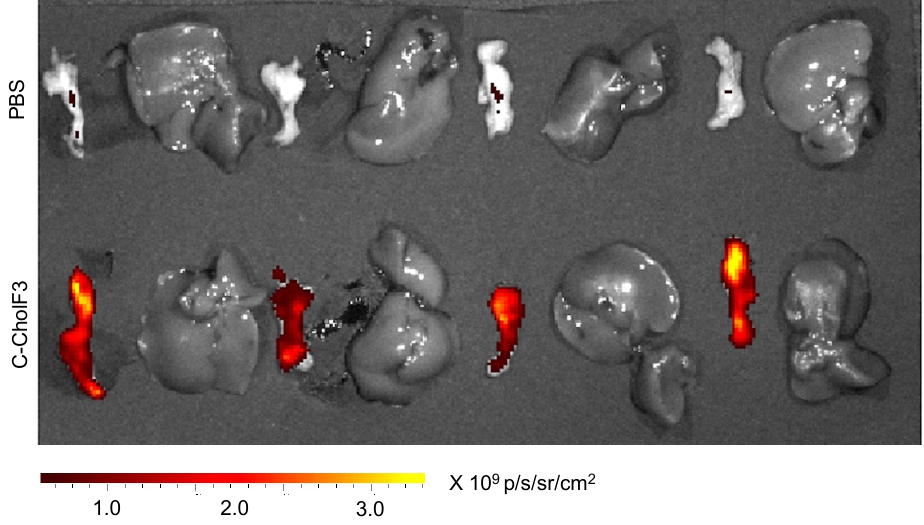

**Figure S13.** Representative tdTomato expression in the pancreas and liver 120h post-injection of C-CholF3 LNPs containing Cre mRNA, and PBS treated control injected intravenously to Ai14 mice at a dose of 1.5 mg/kg (n = 4 biologically independent mice)

**MATERIALS AND METHODS:**

Materials

1,2-di-(9Z-octadecenoyl)-sn-glycero-3-phosphoethanolamine (DOPE), cholesterol and 1,2-dimyristoyl-rac-glycero-3-methoxypolyethylene glycol-2000 (DMG-PEG2K) were purchased from Avanti Polar Lipids, DLin-MC3-DMA, SM-102, 306Oi10, C12-200, Vitamin A, Vitamin D3, Vitamin K1, and (±)-α-Tocopherol was purchased from Cayman Chemicals. Riboflavin and Triton X-100 was purchased from Sigma. THP1 was synthesized in our lab using our established method.^[1]^ Fetal Bovine serum was purchased from Gibco. QuantiT RiboGreen RNA Assay Kit was purchased from Invitrogen. Clean cap FLuc-mRNA and CleanCap Cre mRNA was purchased from Trilink Biotechnology. HEK 293, HUVEC and RAW 264.7 cell lines purchased from ATCC. HFF and HMC3 cell lines were given by Dr. Seungman Park’s lab at UNLV. All the cells were cultured according to the ATCC guidelines. DMEM and MEM growth medium (Gibco, USA) containing sodium bicarbonate, without sodium pyruvate and HEPES, was supplemented with 10% fetal bovine serum (Gibco, USA) and 1% penicillin/streptomycin (Thermo Fisher Scientific). Human umbilical vein endothelial cells were maintained in Ham’s F12K medium (ATCC, USA) supplemented with 10% fetal bovine serum, 1% penicillin-streptomycin-amphotericin B (Fungizone) mix (BioWhittaker, Walkersville, Md.), 30 μg of endothelial cell growth supplement per ml, and 100 μg of heparin (Sigma) per ml.

Formulation of Lipid Nanoparticles

The organic phase was prepared by dissolving the corresponding ionizable Lipid, DOPE, cholesterol, DMG-PEG2K, and the fifth component in ethanol at a molar ratio mentioned in Table S1. The generic four-component LNPs (SM-102, DLin-MC3-DMA, C12-200) which were used as controls were formulated in a molar ratio of 35:16:46.5:2.5. The aqueous phase was prepared by dissolving the corresponding mRNA in 10 mM citrate buffer at pH 3 (Teknova, Hollister, CA, USA). The ionizable lipid to mRNA weight ratio for all LNPs was 10:1.50. The hundred formulations were formulated in a 96-well plate, where each well had the ethanol phase and the aqueous phase was mixed rapidly in the well using a multichannel pipette. During validations, the two phases were loaded into separate glass syringes (Hamilton Company, Reno, NV) and LNPs were formed by chaotic mixing of the organic and aqueous phases at a 1:3 volume ratio in a microfluidic device using Fusion 400 X (Chemyx Inc, USA). The LNPs were subsequently dialyzed against 1X PBS (Thermo Fisher Scientific, Walthman, MA, USA) in 20 kDa molecular weight cutoff dialysis cassettes (Thermo Fisher Scientific) for 4 hours.

Characterization of Lipid Nanoparticles

The encapsulated mRNA concentration and encapsulation efficiency of the LNPs were assessed using the Quant-iT RiboGreen assay (Thermo Fisher Scientific), following established protocols.^[2,3]^ Each LNP sample was diluted 100-fold in two microcentrifuge tubes, one containing 1X TE buffer and the other containing 1% (v/v) Triton X-100 (Sigma, USA) in 1X TE buffer. The Triton X-100 samples were mixed thoroughly and incubated for 5 minutes to lyse the LNPs. Samples (LNPs in 1X TE buffer, LNPs in 1% Triton X-100, and mRNA standards) were placed in quadruplicate in black-walled 96-well plates. The RiboGreen detection reagent was then added to each well according to the manufacturer’s instructions. The plate was shaken at 200 rpm in the dark for 5 minutes, and fluorescence intensity was measured using a GloMax Explorer plate reader (Promega, USA) with an excitation wavelength of 480 nm and an emission wavelength of 520 nm. The encapsulated mRNA concentration was calculated from a standard curve generated using univariate least-squares linear regression. Encapsulation efficiency (EE) was determined using the formula: $1-\frac{R_{TE}}{R_{TX}}$

where RTERTE is the free RNA content in TE buffer, and RTXRTX is the total RNA content in 1% Triton X-100 buffer. The hydrodynamic diameter and polydispersity index (PDI) of the LNPs were measured using a Mobius instrument (Wyatt Technology, Santa Barbara, CA, USA). Each LNP sample was diluted 100-fold in 1X PBS and placed in a cuvette (Wyatt Technology) for analysis. The surface ζ-potential was also measured using the Mobius, with each LNP sample diluted 100-fold in deionized water (Thermo Fisher Scientific) and placed into a capillary cell for measurement. Size and ζ-potential were reported as mean ± standard deviation (n = 3 technical replicates).

TNS Assay

The pKa of LNPs was determined using a TNS (6-(p-Toluidino)-2-naphthalenesulfonic acid) binding assay. A stock solution of 0.16 mM TNS reagent (Sigma Aldrich) was prepared in deionized water. LNPs were diluted to a concentration of 40 ng/mL, and 10 µL of the TNS stock solution was added to each well. The final volume per well was adjusted to 250 µL, consisting of 150 mM sodium chloride, 20 mM sodium phosphate, 20 mM ammonium acetate, and 25 mM ammonium citrate. The assay was conducted across a pH range of 2 to 12, with increments of 0.5 pH units. Samples were placed in black 96-well plates and mixed on a plate shaker at 300 rpm for 5 minutes at room temperature in the dark. Fluorescence measurements were taken using a GloMax Explorer plate reader (Promega), with excitation and emission wavelengths set at 322 nm and 431 nm, respectively, each with a 20 nm bandwidth and a gain of 60. Normalized fluorescence data were plotted against pH, and the pKa value was determined as the pH corresponding to the inflection point of the titration curve.^[4]^

In Vitro Studies

Cells were seeded at a density of 18,000 cells per well in 100 µL of DMEM in a 96-well plate and allowed to adhere for 24 hours. After this, the media was removed, and 75 µL of fresh DMEM without penicillin-streptomycin was added. The cells were then treated with 125 ng of mRNA per 18,000 cells to evaluate in vitro luciferase expression mediated by each LNP. DMEM alone served as the negative control, while LNPs formulated with MC3 were used as the positive control. The LNP-treated cells were incubated at 37°C for 24 hours. Following incubation, 100 µL of luciferase assay substrate (Promega) was added to each well. The plate was shaken on a plate reader at 200 rpm in the dark for 10 minutes, and luminescence intensity was measured using the GloMax Explorer plate reader (Promega, USA).

Confocal Scanning Laser Microscopy (CLSM)

HMC3 cells were seeded in glass bottom dishes (Thermo Fisher, USA) and incubated for 24 h. The medium was then replaced with a culture medium containing EGFP/LNPs (EGFP:125 ng) for transfection at different times (30 mins, 1, 2, 4 hr). Briefly, LNPs encapsulating 125 ng of GFP mRNA LNPs labeled with Did (0.5% molar ratio of LNPs) were added to the dishes. Cell nuclei and lysosomes were stained with Hoescht 33342 (Lumiprobe, USA). Images were obtained using a confocal scanning laser microscope (Nikon A1, Japan) with an oil immersion 40× objective lens. The imaging parameters were kept constant during the experiments.

Tissue Immunohistochemistry (IHC) and H&E Stain

Hematoxylin and eosin (H&E) staining and immunohistochemistry (IHC) on pancreas and liver tissues from C57BL/6J mice were performed on formalin-fixed paraffin-embedded (FFPE) sections. Tissues were post-fixed in 10% formalin for 24–48 hours at room temperature (with the option to remain in formalin for up to a week) and then transferred directly to 70% ethanol. Histological processing was completed by HistoWiz Inc. (NY, USA) following their Standard Operating Procedure and fully automated workflow. Samples were embedded in paraffin and sectioned at 4 µm thickness. IHC was carried out on a Bond Rx autostainer (Leica Biosystems), utilizing heat-mediated antigen retrieval with Epitope Retrieval Solution 1 (Leica Biosystems). Staining was performed using the Bond Polymer Refine Detection kit (Leica Biosystems), adhering to the manufacturer’s protocol. After staining, tissue sections were dehydrated and coverslipped using a TissueTek-Prisma and Coverslipper system (Sakura). Whole slide scanning at 40x magnification was performed using an Aperio AT2 system (Leica Biosystems).

Multiplex Analysis of Cytokines

This study used Luminex xMAP technology for multiplexed quantification of 10 Mouse cytokines, chemokines and growth factors. The multiplexing analysis was performed using the Luminex™ 200 system (Luminex, Austin, TX, USA) by Eve Technologies Corp. (Calgary, Alberta).  Ten markers were simultaneously measured in the samples using Eve Technologies' Mouse Focused 10-Plex Discovery Assay® (MilliporeSigma, Burlington, Massachusetts, USA) according to the manufacturer's protocol.  The 10-plex consisted of GM-CSF, IFNγ, IL-1β, IL-2, IL-4, IL-6, IL-10, IL-12p70, MCP-1, and TNFα.  Assay sensitivities of these markers range from 0.4 – 10.9 pg/mL for the 10-plex.  Individual analyte sensitivity values are available in the MilliporeSigma MILLIPLEX® MAP protocol.

References

[1]	I. Isaac, A. Shaikh, M. Bhatia, Q. Liu, S. Park, C. Bhattacharya, *ACS Nano* **2024**, *18*, 29045.

[2]	A.-G. Reinhart, A. Osterwald, P. Ringler, Y. Leiser, M. E. Lauer, R. E. Martin, C. Ullmer, F. Schumacher, C. Korn, M. Keller, *Mol. Pharmaceutics* **2023**, *20*, 6492.

[3]	L. Cui, S. Pereira, S. Sonzini, S. van Pelt, S. M. Romanelli, L. Liang, D. Ulkoski, V. R. Krishnamurthy, E. Brannigan, C. Brankin, A. S. Desai, *Nanoscale* **2022**, *14*, 1480.

[4]	M. J. Carrasco, S. Alishetty, M.-G. Alameh, H. Said, L. Wright, M. Paige, O. Soliman, D. Weissman, T. E. Cleveland, A. Grishaev, M. D. Buschmann, *Commun Biol* **2021**, *4*, 1.
